# Safety of Non-invasive Brain Stimulation in Patients with Implants: A Computational Study

**DOI:** 10.1101/2024.04.19.590046

**Authors:** Fariba Karimi, Antonino M. Cassarà, Myles Capstick, Niels Kuster, Esra Neufeld

## Abstract

**Objective:** Non-invasive brain stimulation (NIBS) methodologies, such as transcranial electric (tES) and magnetic stimulation are increasingly employed for therapeutic, diagnostic, or research purposes. The concurrent presence of active or passive implants can pose safety risks, affect the NIBS delivery, or generate confounding signals. A systematic investigation is required to understand the interaction mechanisms, quantify exposure, assess safety, and establish guidance for NIBS applications.

**Approach:** We used measurements, simplified generic, and detailed anatomical modeling to: (i) systematically analyze exposure conditions with passive and active implants, considering local field enhancement, exposure dosimetry, tissue heating and neuromodulation, capacitive lead current injection, low-impedance pathways between electrode contacts, and insulation damage; (ii) identify safety metrics and efficient prediction strategies; (iii) quantify these metrics in relevant exposure cases and (iv) identify worst case conditions. Various aspects including implant design, positioning, scar tissue formation, anisotropy, and frequency were investigated.

**Results:** At typical tES frequencies, local enhancement of dosimetric exposure quantities can reach up to one order of magnitude for DBS and SEEG implants (more for elongated passive implants), potentially resulting in unwanted neuromodulation that can confound results but is still 2-3 orders of magnitude lower than active DBS. Under worst-case conditions, capacitive current injection in the lead of active implants can produce local exposures of similar magnitude as the passive field enhancement, while capacitive pathways between contacts are negligible. Above 10 kHz, applied current magnitudes increase, necessitating consideration of tissue heating. Furthermore, capacitive effects become more prominent, leading to current injection that can reach DBS-like levels. Adverse effects from abandoned/damaged leads in direct electrode vicinity cannot be excluded.

**Significance:** Safety related concerns of tES application in the presence of implants are systematically identified and explored, resulting in specific and quantitative guidance and establishing a basis for safety standards. Furthermore, several methods for reducing risks are suggested.

## 1. Introduction

Disorders impacting brain function, such as stroke, epilepsy, Parkinson’s and Alzheimer’s disease, alongside mental health conditions like schizophrenia and depression, pose significant and escalating challenges to societal well-being [1]. Such diseases often lack satisfactory treatment options with current therapeutic approaches. Over the past few decades, substantial research efforts have been dedicated to developing treatments and cures for these disorders with the aim to improving the patient’s quality of life. Non-invasive brain stimulation (NIBS) has emerged as a promising technique for a wide range of conditions, including those for which conventional pharmacological approaches are ineffective [2]. These methods produce minimal adverse effects and avoid invasive procedures and, when personalized for precise targeting, NIBS can provide a safe and effective therapeutic solution. Transcranial electric stimulation (tES) uses electric currents applied to the scalp while transcranial magnetic stimulation (TMS) utilizes magnetically induced currents in the brain to modulate neural activity at a range of spatial and temporal scales through electromagnetic (EM) coupling with neural electrophysiology. tES, including transcranial alternating current stimulation (tACS) and transcranial direct current stimulation (tDCS), typically induces subthreshold changes in transmembrane potentials of cortical neurons [3]. For an extended discussion of interaction mechanisms and associated exposure quantities-of-interest, see [4]. Electrode shape and placement, along with stimulation parameters, such as amplitude, duration, and pulse shape, determine the exposure conditions (current density, charge accumulation, etc.). tDCS primarily involves anode-cathode pairs to modulate neuronal excitability [5]. tACS, on the other hand, entrains brain oscillations at specific frequencies, impacting cognitive or other brain functions [6]. NIBS techniques, such as tES and TMS, have high potential for a diverse range of clinical applications [7, 8]. Recently, temporal interference stimulation (TIS) [9], has been proposed that facilitate the non-invasive stimulation of deep targets by leveraging multiple applied electric (E-)fields. These field operate at frequencies too high to induce typical neuromodulation at conventional current strengths due to the neural low-pass membrane filtering. However, the difference in their frequencies within the physiological range allows them to impact neural activity by modulating field amplitude and orientation where currents overlap. While the detailed mechanisms of low frequency modulated kHz fields are still unclear [4], first applications are already showing intriguing and promising results [10, 11, 12]. In contrast to tES, TMS, which has been applied since 1985 [13], induces electrical currents in the brain through a time-varying magnetic field generated by passing high currents through one or multiple insulated coils on or near the scalp [14]. TMS has been found to be an effective therapy for a range of neurological disorders, including Parkinson’s disease, Alzheimer’s disease, depression, attention deficit and hyperactivity disorders, dyslexia, autism spectrum disorders, to name but a few [7].

The increasing use of NIBS as both a research and therapeutic tool is accompanied by an increasing number of patients with implants. In this patient population, electrodes for deep brain stimulation (DBS) and recording (stereoelectroencephalography, SEEG) are frequently used, which raises significant concerns about the safety of employing NIBS in the presence of metallic implants. Typical safety recommendation for tES and TMS advise exclusion of subjects with active or passive implanted electrodes [15, 16]. This precaution aims to prevent potential adverse effects (typically thermal tissue damage or unwanted stimulation of neighbouring areas), leading to the exclusion of patient groups that are in need for advanced therapies. Thus, a good understanding combined with guidance are crucial to determine under which conditions administration of NIBS is still safe or how it should be adapted to ensure safety.

Few studies have reported on the safety of combining *tES with DBS and SEEG*. In one instance, cumulative tDCS over the primary motor cortex was explored as a potential therapy to alleviate freezing of gait in Parkinson’s disease patients with DBS without observing significant adverse effects [17]. However, only a small sample size (two patients) was considered and no systematic investigation was performed, which limits the generalizability of the findings. The same is true for a case study in a single female patient with implanted DBS electrodes, where tDCS was used as an adjuvant therapy for the treatment of generalized dystonia and proved to be well tolerated [18]. In addition to limited sample size, the absence of long-term follow-up data prevents assessment of sustained effects and potential risks associated with such a combination therapy [18]. Comparisons of various tDCS montages, both with intact skulls and simulated burr holes, indicated little effect on peak current density and E-fields, suggesting that the presence of a DBS lead might not significantly alter brain current flow under realistic conditions when the burrhole defect is fluid-filled and relatively conductive [19]. However, it is important to note that these simulations did not account for the local field enhancement caused by metallic contacts in the lead and the metallic guidance screw for DBS leads, which could substantially increase peak density and E-field. The authors conclude that, while theoretical concerns exist, current evidence, derived from limited theoretical and clinical experience, does not conclusively support an elevated risk of serious adverse effects associated with implants during tDCS. In view of the small number of published investigations (see citations above), the anecdotal nature of much of the *in vivo* evidence, and the lack of systematic approaches, further research is needed to establish reliable safety guidelines for the application of NIBS in patients with active and passive implants.

More data are available on the safety of TMS in the presence of metallic implants. The following briefly reviews the latest systematic studies on the safety of TMS with implants (focusing on DBS and SEEG):

There are conflicting results on *TMS alongside DBS*. Some studies indicate that no significant E-field enhancement results [20], while others report unsafe induced currents in the presence of implanted DBS probes [21, 22]. Shimojima et al. [23] investigated the safety of TMS in DBS patients using a plastic phantom filled with tissue simulating media. They found no mechanical movement or temperature increase after one hour of 0.2 Hz repetitive TMS (rTMS) delivered using a Magstim 200 stimulator (The Magstim Co., Ltd, Dyfed, UK) operating at 100 % intensity. However, they observed that TMS intensities over 50 % led to evoked charge injection exceeding 30*µ* C*/*cm^2^ per phase – the limit recommended by [24] to prevent tissue damage – in addition to posing the risk of unwanted neurostimulation near the implant. These risks further increase when DBS leads are looped, as induced lead currents become more relevant. While the study suggests that TMS can be used safely in patients with DBS implants, provided that the coil placement is chosen to minimize implant exposure, several important limitations must be considered: (i) only one specific DBS lead was investigated, focusing on the magnitude of the induced current without considering the spatial current density and E-field distribution, (ii) a simplified homogeneous phantom of limited realism was employed, and (iii) the DBS location was not varied. Mhaskar et al. [25] employed finite element modeling (FEM) to assess EM interference and E-field distributions in DBS patients undergoing TMS and concluded that this can be done safely under certain conditions (e.g., avoiding direct targeting of DBS lead windings, using focal TMS). However, also this study has important limitations as only one DBS lead and implant design and two implantation locations were considered (the TMS coil location and orientation were varied), and the E-field enhancement was determined only in gray matter, without consideration for local field enhancement near metallic contacts.

Wang et al. explored *TMS in combination with SEEG*, revealing a favorable safety profile with no reported adverse events [26]. However, this study also focused on a single SEEG implant design, employed a simple phantom, and only measured induced voltages without measuring or computing exposure quantities-of-interest such as local field enhancement near metallic contacts, which could result in unwanted neurostimulation. Moreover, the location of the TMS coil and orientation was not varied. While the reported findings on TMS and SEEG are mostly reassuring, they remain fragmentary and are based on a small number of considered cases. In view of the important investigative, diagnostic, and therapeutic benefits that TMS-SEEG could offer, more systematic safety investigations are desirable.

Golestanirad et al. assessed safety concerns related to Lorenz forces induced by *TMS in electrocorticography* (ECoG) arrays and resulting shear stress in the brain using FEM modeling [27]. They found that the induced stress in brain tissue is below the limit of tissue damage. However, that study exclusively focused on Lorenz-forces-related risks, which is only one of the potentially adverse biophysical interactions. In another study, Varnerin et al. investigated *TMS in the presence of metallic stents* and did not find significant heating for common rTMS protocols for specific stent types [28]. However, the study is limited in as they used simplified phantom conditions and only investigated a small number of coil geometries and orientations.

Although tES applications date back to 1874 [29], comprehensive *guidelines and safety regulations* only started appearing recently. In 2016, Bikson et al. [19] integrated evidence from human trials, animal models, and computational modeling on tDCS safety. Notably, in the context of subjects with implants, the study states: *“While pre-existing implants remain a theoretical concern, neither theory nor limited clinical experience establish evidence for increased risk of Serious Adverse Effects”*. In 2017, Antal et al. [30] compiled tES safety recommendations based on expert interviews and an extensive literature review. Their work provides fundamental guidance regarding the safety of applying tES to human subjects, the reporting of tES experiments in research and clinical practice, and regulatory issues. Regarding tES in subjects with intracranial implants, they state that *“tES should be applied in subjects with implants only in well-supervised and controlled studies”*. However, clear criteria are not provided. The recent review of Bikson et al. [31], offers guidance for limited-output tES. They aim to minimize risk without limiting the adoption of innovative tES-based technologies and suggest that issuing general warnings or restrictions may not be justified considering the absence of observed severe adverse effects. Conversely, it discourages relying solely on the existing history of safe use as conclusive evidence. [31]underscores the importance of quantitative and scientific risk assessments. Therefore, there is a need for systematic assessment of tES safety for subjects with implants.

The *present study* focuses on safety of tES in the presence of both active (primarily DBS and SEEG) and passive (e.g., stents) implants. The less controlled situations when implants are abandoned (cut) or damaged (see Fig. 1) are also analyzed and discussed. It is concerned with high energy deposition and induced brain heating, unwanted neurostimulation, deviation of tES currents from their intended targets, and the confounding of neural recordings. Quantitative conclusions are drawn with regard to established exposure limits or by relative comparison with common brain stimulation paradigms. This study considers the following mechanisms to be of primary relevance when exposing patients with implanted electrodes (see Fig. 2):

*•* localized, but potentially strong, E-field enhancement due to the presence of highly conductive (e.g., metallic) contacts that concentrate the field lines; if sufficiently strong, enhancement can lead to induced heating [32] or unwanted stimulation (the latter right next to where invasive electrodes might record activity, confounding corresponding measurements)
*•* capacitive coupling between wires in the lead, resulting in lower impedance pathways between electrode contacts that could result in field enhancements or distort the intended brain exposure and perturb stimulation targeting
*•* capacitive current injection into the implant leads (despite intact insulation) where tES electrodes are in close proximity to the lead routing (e.g., proximity to subcutaneous coiling of DBS leads), which could produce high fields and power deposition near implant contacts

**Figure 1.**
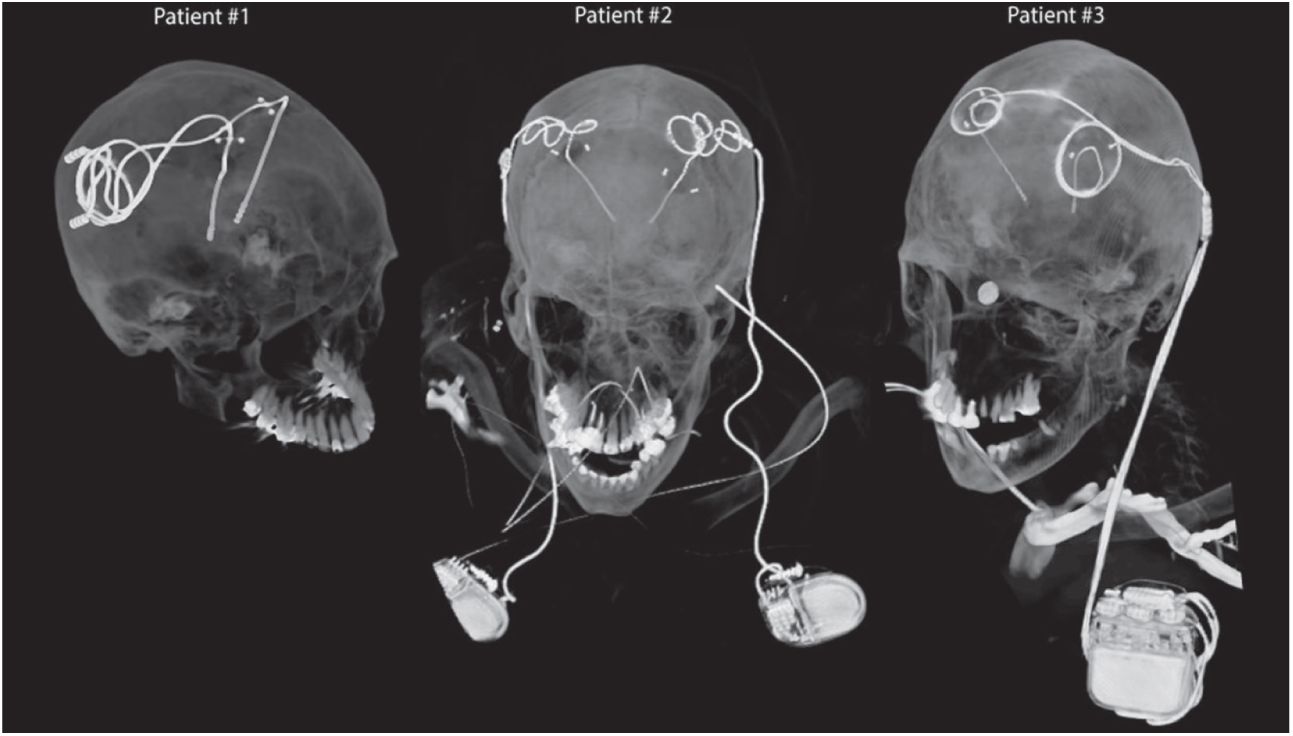
Computed tomography (CT) images obtained postoperatively from patients with isolated and fully implanted DBS systems (reprinted with permission from [33]).

**Figure 2.**
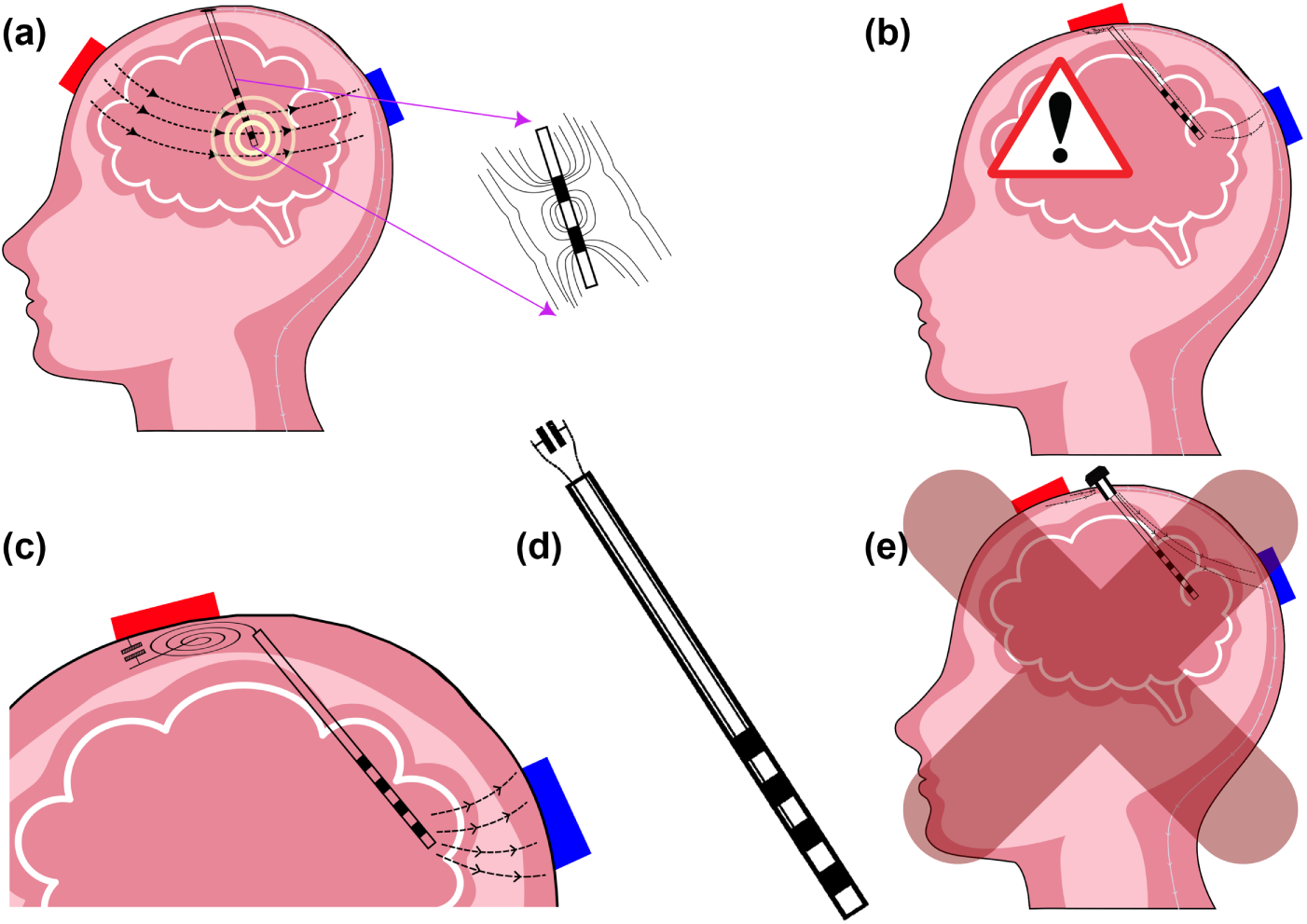
Schematic representations of (a) field-lines and local field enhancement near metallic structures during NIBS; (b) NIBS application in the presence of abandoned leads or leads with damaged insulation; (c) capacitive current injection into a lead passing/coiling near a NIBS electrode; (d) capacitive coupling between lead wires resulting in low inter-contact impedance; (e) NIBS application in the presence of burr holes, which could constitute a risk, but is not covered in the scope of this study.

Another important concern are low impedance pathways for the applied tES currents through burr holes or fixation screws used for implant insertion. Such pathways can result in increased brain exposure, as well as deviations from the intended stimulation targeting. However, systematic treatment of such effects is beyond the scope of this study.

In summary, the goals of this paper are to i) systematically identify and explore safety related concerns of tES application in the presence of implants (with a primary focus on DBS and SEEG implants), ii) establish (where possible quantitative) guidance for tES exposure work, potentially in preparation for safety guidelines and standards, and iii) to identify aspects that require further investigation.

## 2. Methods

The different concerns listed in Sec. 1 were addressed as follows:

*• Localized field enhancements near highly conductive structures:* Systematic exploration (Sec. 2.2.2) of the SEEG and DBS implant geometry parameter space, as well as generic elongated implants, in a simplified simulation setup to determine local enhancement factors for the different safety-relevant quantities-of-interest (QoIs; see Sec. 2.1). In addition, simulations in a detailed anatomical model (Sec. 2.2.5) were used to confirm that the enhancement factors from the simplified model remain applicable.
*• Low impedance between implant contacts:* In view of the variability in implant designs, such impedances are best characterized experimentally. Corresponding illustrative measurements were performed for representative DBS and SEEG implants (Sec. 2.4).
*• Capacitive coupling into implant leads:* As the variability of lead designs favors experimental characterization, a series of measurements have been performed to assess the magnitude of this effect (Sec. 2.4).
*• Ohmic coupling into implant leads:* Direct contact of the lead wires with the tissue when the lead is abandoned (cut) or when the insulation is damaged (Sec. 2.6).

### 2.1. Exposure QoIs

The following dosimetric QoIs were extracted:

- E_ICNIRP,n_: volume-averaged vectorial E-field enhancement factor; ratio of the peak of the volume-averaged vectorial E-field and the incident field in the absence of the implant; averaging followed the ICNIRP 2010 [34] procedure with a sliding cube of 0.1 mm side length instead of 2 mm;
- E_IEEE,n_: line-averaged vectorial E-field enhancement factor; ratio of peak of the line-averaged vectorial E-field and the incident field in the absence of the implant; averaging followed the IEEE 2005 [35] procedure with a straight line of 0.1 mm instead of 5 mm;
- psSAR_0.1g,n_: psSAR enhancement factor; ratio of the peak spatial average specific absorption rate (psSAR) and the reference SAR in the absence of the implant; as commonly employed for dosimetric safety assessment; averaging followed the IEEE 1528 standard [36] procedure with a cube of 0.1 g instead of 10 g;
- *• G* = *max*(*|Eigv*(*H*(*x, y, z*)*|*): spatial peak of maximal absolute Eigenvalue of the Hessian *H*(*x, y, z*) of the electric potential *EP* (*x, y, z*) near the implant; it computes the strongest E-field gradient and is the 3D equivalent of the Activation Function (AF) [37], a common predictor of axonal neurostimulation; the associated Eigenvector corresponds to the direction of that worst-case gradient.

The 0.1 mm averaging length was chosen as a compromise between the length scale of the rapid field decay near metallic contacts and the length-scale of internodal separation of myelinated axons (0.1-2 mm; the internodal separation results in a depolarization proportional to the finite difference approximation of the electric potential gradient, such that the correspondingly averaged E-field correlates maximally with neuromodulation impact). In addition, the influence of increasing that length up to 0.2 mm was studied, but was found to have little impact on the behavior of the local field enhancement factor. The 0.1 g averaging mass was chosen, as it maximizes correlation with induced heating for highly localized exposures.

In addition, the thermal exposure was quantified as:

- ΔT_max,n_: peak temperature increase normalized to the reference temperature in the absence of the implant; no averaging is required because of thermal diffusion.

All QoIs were expressed relative to the EM exposure, respectively heating, in the absence of the implant denoted by subscript *n*.

### 2.2. EM Exposure

All simulations were performed using the Sim4Life platform V7.2 for computational life sciences (ZMT Zurich MedTech AG, Switzerland).

To determine EM fields, EM-exposure QoIs, and sources for the simulation of induced heating, FEM simulations were carried out with the low frequency (LF) solver in Sim4Life. In view of the stimulation frequency range of interest (1 Hz – 100 kHz; the upper limit relates to recent interest in high frequency TIS) the ohmic-current-dominated electro-quasistatic (EQS) approximation is suitable [38], which solves the Laplace equation *∇ ·* (*σ∇ϕ*) = 0. Here *σ* and *ϕ* are the electric conductivity distribution and a scalar field potential, respectively, and the E-field and current density are obtained as **E** = *−∇ϕ* and **j** = *σ***E**. This equation approximates the full Maxwell equations well, as long as ohmic currents dominate over displacement currents and characteristic exposure lengths are short compared to the wavelength. A rectilinear adaptive discretization was employed and a grid-convergence studies was performed (Sec. 2.2.4). The strictest solver convergence criterion (reduction of the residuals by twelve orders of magnitude) was used to guarantee convergence. Metallic parts were treated as perfect electric conductors (PEC) and insulting parts were assigned zero conductivity.

#### 2.2.1. Generic Implant Geometries

A generic, parameterized implant geometry was generated that consisted of three cylindrical contacts (the last one capped by a semi-sphere) interspaced by insulating cylinders, and on top of a cylindrical insulating shaft (see Fig. 3(a). The geometry parameters were varied as follows:

*• d*: implant diameter (0.8, 1.3, and 1.4 mm);
*• L_c_*: contact length (0.5, 1.0, 1.5, 2.0, 2.5, and 3.0 mm);
*• L_i_*: insulation length (0.5, 1.0, 1.5, 2.0, and 4.0 mm).

**Figure 3.**
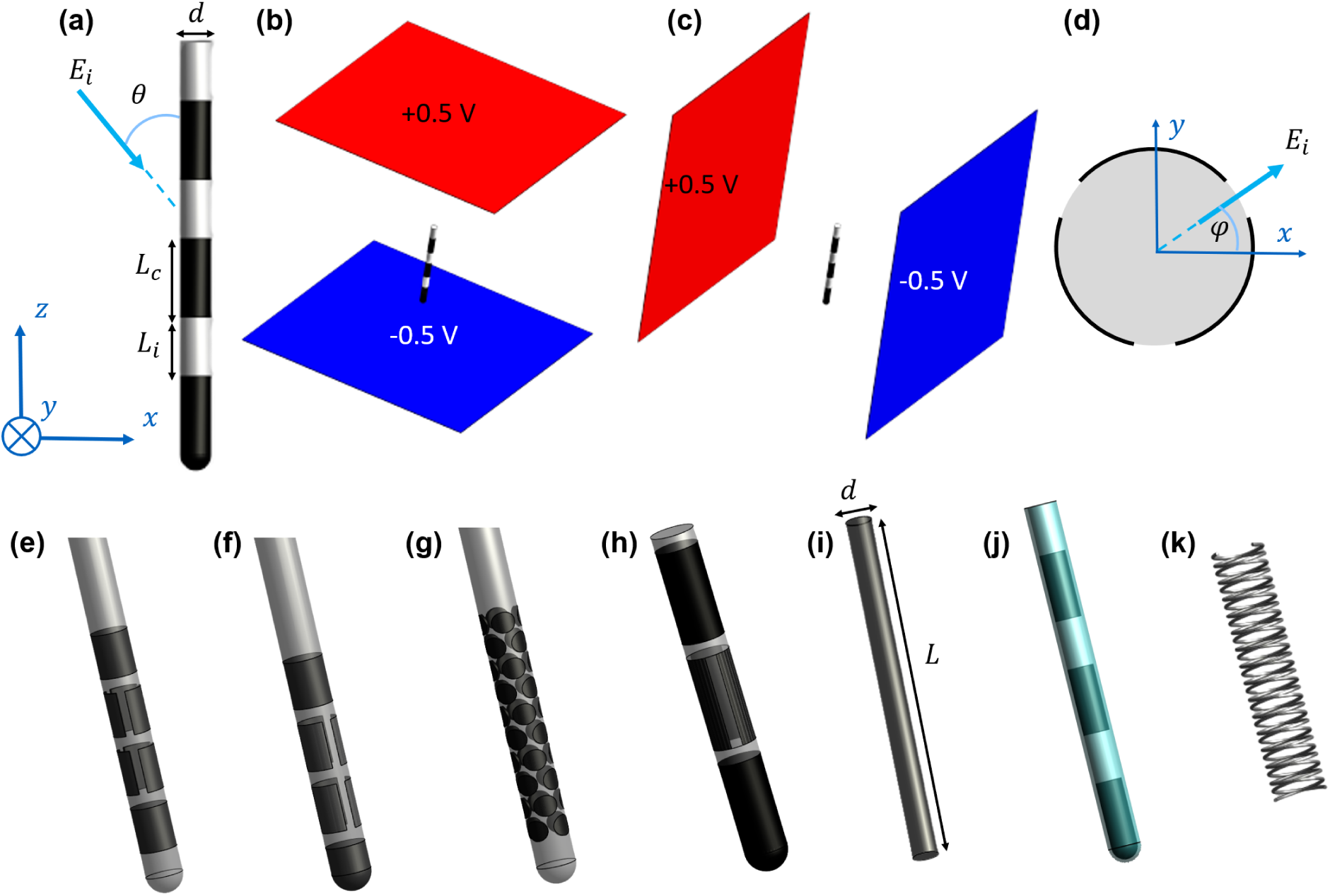
(a) Generic implant geometry used to study local field enhancement (*E_i_*: incident E-field, *θ*: angle of incidence, d: diameter, *L_c_*: contact length, *L_i_*: insulation length), (b) parallel field exposure configuration, and (c) perpendicular configuration. (d) Representation of E-field angle of incidence in the transversal plane for segmented contacts. (e) Abbott/St. Jude directed 6172 DBS lead model (*d*=1.29 mm, *L_c_*=1.5 mm, and *L_i_*=0.5 mm). (f) Boston Scientific Cartesia DBS lead model (*d*=1.3 mm, *L_c_*= 1.5 mm, and *L_i_*=0.5 mm). (g) Medtronic SureStim DBS lead model (*d*=1.27 mm, 40 individual stimulation points with 0.66 mm diameter). (h) Generic implant geometry with segmented contacts, similar to Abbott/St. Jude directed 6172 and Boston Scientific Cartesia. (i) Geometry of elongated simple implant. (j) Generic implant surrounded by scar tissue. (k) Typical stent model (*d*=3.0 mm, *L*=15.0 mm, distance between turns=1.0 mm, wire diameter=0.1 mm)

The ranges were chosen to comprehensively cover commercially available SEEG [39, 40] and DBS implants [41]. It was computationally ascertained that having more than three contacts did not significantly alter the obtained exposure enhancements.

In addition, *segmented electrodes* modelled on Abbott/St. Jude directed 6172 and Boston Scientific Cartesia DBS leads were studied (Fig. 3(e,f)) [41], and *implants with small, circular-shaped contacts (high-density ‘HD’ lead)*, as present in the Medtronic SureStim device, were modelled (Fig. 3(g))[42]. Finally *elongated conductive cylinders* were simulated to mimic the presence of elongated passive implants, such as stents (Fig. 3(i)). For this geometry, the diameter and the length were varied from 1 to 5 mm (in steps of 1 mm) and from 5 to 50 mm (in steps of 5 mm), respectively. This is in alignment with currently available intracranial stents which range in diameter from 2 to 6 mm and in length from 4.2 to 40 mm [43, 44, 45, 46, 47, 48, 49, 50, 51]. For comparison, a detailed geometry from a typical stent was simulated (see Fig. 3(k); stent diameter: 3 mm; length: 15 mm, wire diameter: 0.2 mm, distance between turns: 1 mm) and E_IEEE,n_ was extracted (the most conservative QoI).

#### 2.2.2. Simplified Setup

As mentioned in Sec. 1, the presence of a medical implant with conductive contacts gives rise to localized E-field enhancement near exposed metallic parts of both active and passive implants. This enhancement might result in problematic tissue heating and/or unwanted stimulation in the vicinity of the implant. The dominance of the EQS regime suggest that a local field enhancement factor in the brain can be determined, that is primarily influenced by the implant geometry (metallic and insulating parts) and only weakly depends on the details of the anatomical environment. To quantify these enhancement factors, EM simulation were conducted for a simplified homogeneously conductive cubic region. The conductivity was chosen as 0.3 S/m to conservatively represent brain tissue. The phantom dimensions (i.e., 31.5 mm in all directions) were modeled to be 50 % larger than the maximal implant length.

As a consequence of the implant symmetry, any homogeneous exposure condition can be derived from two simulated perpendicular field orientations (see Fig. 3(b,c)). (Averaged) E-fields are easily obtained for any incident angle *θ* (Fig. 3(a)) through weighted superposition. SAR and temperature increase can be computed using a Hermitean quadratic normal form (four independent thermal simulations are required, see [52, 53]). For segmented and HD electrodes, where symmetry is reduced, three simulations with perpendicular fields are needed to recreate any E-field exposure orientation (parameterized in terms of *ϕ* and *θ*; see Fig. 3(a,d)).

The computational domain in the vicinity of the implant, where peak exposures are localized, was discretized with a rectilinear grid of 0.015 mm resolution. This ensures proper handling of the strong field gradients and high dielectric contrast in that region. The convergence analysis revealed that a 1 mm resolution is sufficient elsewhere.

#### 2.2.3. Scar Tissue Encapsulation

Foreign body response frequently results in scar tissue formation during chronic electrode implantation. The scar tissue was modelled as an implant- surrounding layer with a thickness ranging from 25 to 300 *µ*m (25, 50, 75, 100, 200, 300 *µ*m; see Fig. 3(j)); [54] reports thicknesses up to 300 *µ*m for active and 100 *µ*m for passive electrodes. The dielectric properties were based on the mice measurements from [54], which were reported to be *σ* =0.019 S/m at 20 Hz, 0.05 S/m at 1 kHz, and 0.28 S/m at 300 kHz. Simulations were performed with both 0.019 S/m and 0.05 S/m as the conductivity for scar tissue. The 100 kHz condition was considered less relevant for tES, and furthermore, a value of 0.28 S/m would make the scar tissue similar to the simulated brain medium, resulting in behavior resembling the scarless case.

#### 2.2.4. Convergence Analysis

For the convergence analysis, the resolution of the smallest implant (*d*=0.8 mm, *L_c_* = *L_i_* = 0.5 mm; highest discretization sensitivity) was varied in three steps: 0.01, 0.015, and 0.02 mm. At each resolution, E_ICNIRP,n_, E_IEEE,n_, and psSAR_0.1g,n_ were computed for normal and parallel incidence. The discretization error was quantified in comparison to asymptotic (infinitely-fine resolution) reference values, that were estimated by linearly fitting log-log plots of the deviations versus resolution. The resulting bidirectional inter-dependence between asymptotic reference values and fitting is resolved by optimizing the former such that in the latter the regression slopes are close to the order of the numerical scheme and the correlations *R*^2^ are close to 1 (i.e., the discretization error follows a power-law with reasonable exponent). The relative difference of a QoI at resolution *i*, with regard to its asymptotic value at infinite resolution was calculated as 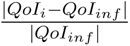 .

#### 2.2.5. Anatomical Model Validation of Enhancement Factor Approach

To verify the applicability of enhancement factors determined under simplified homogeneous exposure conditions, additional simulations with a detailed realistic anatomically human head model derived from multi-model image data were performed. The IXI025 model [55] was used, which includes 15 different tissue classes with isotropic material properties assigned according to the IT’IS LF V4.1 tissue database [56]. Simulations were performed using the Sim4Life LF EM solver. Two tDCS setups were simulated, each comprising a pair of skin surface electrodes (see Fig. 4(a) and Fig. 4(d)) and seven SEEG catheters (*d*=0.8 mm, *L_c_*=2 mm, *L_i_*=1.5 mm, each with 8 to 18 contacts) implanted in the brain, mimicking a real application scenario (see Fig. 4). The SEEG models were inserted in the brain in accordance with real trajectories extracted from image data of an epilepsy patient.

**Figure 4.**
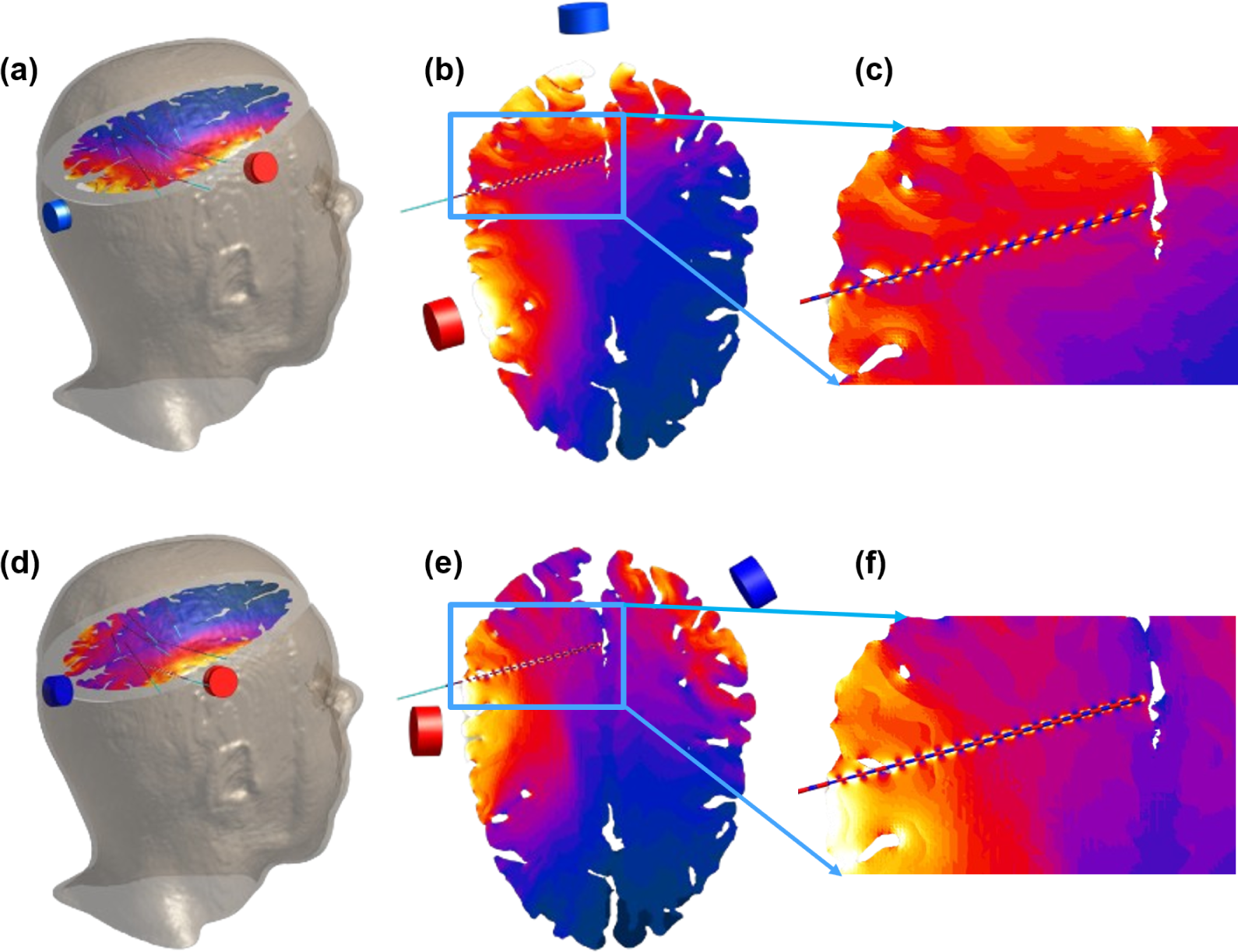
(a) Illustration of the IXI025 model, with tDCS montage 1 and the implanted SEEG electrodes. (b) E-field distribution on a slice containing an SEEG electrode, and (c) zoomed E-field distribution near an SEEG implant. (d, e, f) depict the same as (a, b, c) for tDCS montage 2.

**Figure 5.**
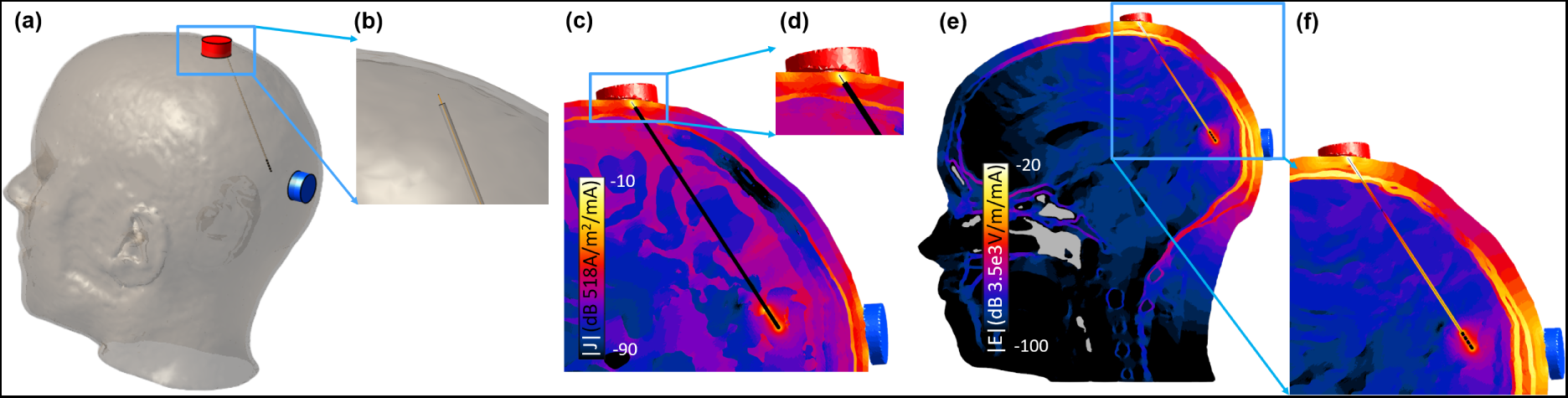
(a,b) Illustration of the IXI025 model with a tDCS montage and an abandoned lead with 0.2 mm diamater lead, along with a zoomed version of the lead tip. (c,d) Current density distribution on a slice containing the lead and zoomed version at the tip. (e,f) E-field distribution on a slice containing the lead and zoomed version near the lead.

EM simulations were conducted both with and without incorporating the SEEG models, utilizing an unstructured mesh with 123 million tetrahedral elements. The target edge length for the implant region and its surroundings was set to 0.2 mm, while a 1 mm target edge length was specified for the rest of the simulation domain.

To enable application of the local enhancement factors, the E-field vector from the exposure simulation without implants was averaged over the surface of each SEEG contact electrode. The magnitude and angle of these averaged E-field vectors relative to the corresponding catheter axes were extracted. Subsequently, they were used to predict the expected local E-field peaks in the presence of the implant, employing the angle-dependent *E_IEEE,n_* enhancement factors from the simplified homogeneous setup (see Sec. 2.2.1). The obtained results were then compared against the local field peaks extracted in the detailed anatomical simulation which included the SEEG implants.

### 2.3. Tissue Heating

The Sim4Life thermal solvers were used to assess EM-induced tissue heating based on the Pennes Bioheat Equation (PBE, [57]):

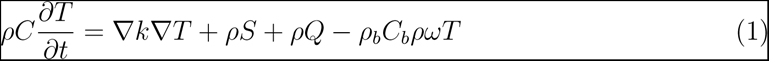

(*ρ*: density, 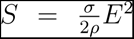: SAR obtained from the EM simulation, *Q*: metabolic heat generation rate, *C*: heat capacity, *ω*: perfusion, *k*: thermal conductivity, *T* : temperature increase; subscript *_b_* denotes a blood property). By focusing on temperature increase, blood temperature, metabolic heating, the heterogeneous initial tissue temperature, and the ambient temperature become irrelevant (provided temperature increase remains below two celcius degrees [58]). Tissue properties were obtained based on brain tissue values from [56]. Conservative insulating (zero-flux Neumann) boundary conditions were applied. Thermal simulations were first conducted with the transient solver for selected cases to show that a steady-state is reliably achieved after 30 minutes of simulated heating. The steady-state thus represents a realistic worst-case scenario. Since simulations with the steady-state solver are computationally much more efficient than simulations with the transient solver, subsequent thermal investigations for the generic implant setup discussed in Sec.2.2.1 were performed with the steady-state solver. These simulations were executed with a maximum of 1000 iterations to achieve a relative residuum convergence below 1*×*10*^−^*^7^.

As mentioned in Sec. 2.2.2, the temperature increase is determined by evaluating a Hermitean quadratic normal form derived from four independent thermal simulations (three when the E-fields are real-valued, as is the case in the EQS approximation). The EM power deposition (heat source) is:

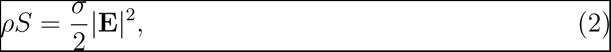

where:

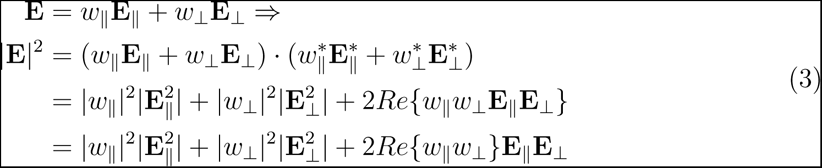

where **E**_∥_ and **E***_⊥_* are implant-axis-parallel and -normal E-field vectors from which **E** can be obtained as a linear combination with weights *w_∥_* = *cos*(*θ*)*|***E***|/|***E***_∥_|* = **E** *·* **E***_∥_/|***E***_∥_|*^2^ and *w_⊥_* = *sin*(*θ*)*|***E***|* = **E** *·* **E***_⊥_/|***E***_⊥_|*^2^. They were arbitrarily set to have 1 V/m magnitude in our simulations. The last equality in (3) follows from the real-valued nature of **E***_∥_* and **E***_⊥_* in the EQS approximation. Because of the linearity of the PBE, three thermal simulations, with heat sources 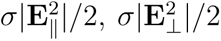, and *σ***E***_∥_***E***_⊥_/*2, are sufficient per implant to derive the temperature increase for any exposure configuration. If the associated temperature increase distribution are denoted as *T_∥_*, *T_⊥_*, and *T_∥⊥_*, the temperature increase resulting from exposure incidence at an angle *θ* can be expressed as:

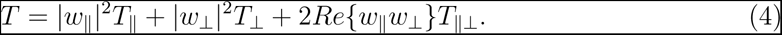

### 2.4. Impedance Measurements

Lead capacitance measurements were carried out using a BK Precision 880 LCR meter for both an AD-Tech macro-micro depth sensing electrode (*d*=1.28 mm, *L_c_*=1.57 mm, *L_i_*=3.43 mm, number of contacts: 6 and 10, lead length *≃* 40 cm from electrode to contact) and a Medtronic SenSight directional lead (B33005 lead with 1-3-3-1 electrode configuration, *d*=1.36 mm, *L_c_*=1.5 mm, *L_i_*=0.5 mm). The Medtronic SenSight directional lead is mostly used for DBS and consists of an implanted part and an extension lead with six stimulation contacts, two ring electrodes (proximal and distal) and two central pairs of electrodes split into two sectors on either side of the lead. Both inter-electrode and lead-to-tissue capacitances were measured. The former are dominated by the capacitances between the individual lead wires connected to the different contacts. They were measured in the absence of tissue, either between electrode contacts or between lead connector contacts – whichever was easiest from a measurement perspective. The capacitance between a lead and its surrounding was measured by closely wrapping the lead in a conductive foil and measuring the capacitance between the foil and a contact. These measurements do not consider contributions from any device to which the lead might be connected, which must be assessed separately.

The relevance of the inter-electrode capacitance can be assessed by comparing it with the corresponding impedance of the ohmic pathway through brain tissue. The latter was determined by simulating a voltage difference between the corresponding contacts in the simplified setup using the Sim4Life EM LF solver and integrating the obtained current density over a surface surrounding one of the electrodes.

To assess the relevance of the capacitive coupling into the lead, the worst-case scenario is considered, in which the lead is mostly coiled below one of the tES electrodes and the induced current exits through an implant contact in the brain near the other tES electrode. Given a capacitance per unit length of *c*, the induced current can be computed as:

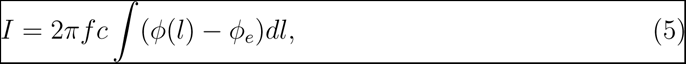

where *f* is the frequency, *ϕ*(*l*) is the incident potential along the lead trajectory and *ϕ_e_* is the potential at the contact. In the above discussed worst-case, the equation reduces to 2*πfcLV* , where *L* is the lead length and *V* is the applied tES voltage. For an implant situated in the brain, this estimation is overly conservative, as the skull barrier will already reduce the voltage difference by 20-40 %.

### 2.5. Neurostimulation

#### 2.5.1. Field Gradient Maximum

To investigate the stimulation potential of local E-field enhancement near implants, the maximal field gradient magnitude *G* was determined (see Sec. 2.1). This QoI was extracted for all generic implants (see Sec. 2.2.1) and field-incidence angle.

#### 2.5.2. Comparison with DBS Activation

To quantitatively relate the stimulation potential of passive implants to that of DBS-like active implants, we compared the magnitude scales of the distributions of the above-mentioned field gradient *G*. This comparison is meaningful because the *G*-distribution of the local field enhancement near a passively exposed electrode and that of a similarly-shaped active electrode (i) display similarly shaped field peaks near electrode edges and (ii) the cumulative dose-volume histograms (DVH) of the maximal field gradient distributions are nearly identical, up to a constant scale factor (see Fig. 15(d)). This scale factor is determined by comparing the enhancement E-field distribution resulting from passive exposure to a 1 V/m incident E-field to the E-field distribution generated by a monopolar cathodic stimulation with 1 mA injected current (modeled by applying isopotential Dirichlet conditions to all the contacts while keeping the sufficiently distant domain boundaries at ground). In other words, the scaling factor leading to a comparable volume of tissue activation (VTA; a common metric in DBS impact analysis [59]) is determined. Note, that while the E-field peak shape and DVH are similar, the field polarity is not. For example, passive exposure to a parallel field leads to regions of local E-field enhancement and regions of local E-field reduction in proximity of the lead, therefore of both polarities. In the case of active stimulation, the E-field is always monopolar, with the polarity depending on the type of electrode polarization (cathodic or anodic).

### 2.6. Abandoned or Damaged Leads

Abandoned (cut) or leads with damaged insulation close to a NIBS electrode provide a direct current pathway into the brain. This could potentially lead to DBS-like currents exciting from implant contacts. To conservatively investigate that case, a simulation featuring a detailed head model and a cut lead with a protruding wire (modelled with 0.2 and 1 mm diameter) connected to the implant contacts was simulated (see Fig. 5). The fraction of applied tES current exiting through the implant contacts is 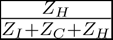, where *Z_H_* is the head impedance between the tES electrodes, *Z_C_* is the impedance for entering the cut/damage location, and *Z_I_* is the impedance between the implant contacts and the return electrode.

## 3. Results

### 3.1. Convergence Analysis

The results of the generic setup convergence analysis for E_ICNIRP,n_, E_IEEE,n_, and psSAR_0.1g,n_ enhancement factors are presented in Fig. 6 and Fig. A.1 for perpendicular and parallel incidence, respectively. The psSAR_0.1g,n_ discretization errors are so small that they reach the noise level (hence the reduced *R*^2^ and meaningless slope). Based on these results, a grid resolution of 0.015 mm was selected, which produces deviations that remain below 10 % for all QoIs (except for E_IEEE,n_ in the case of parallel incidence).

**Figure 6.**
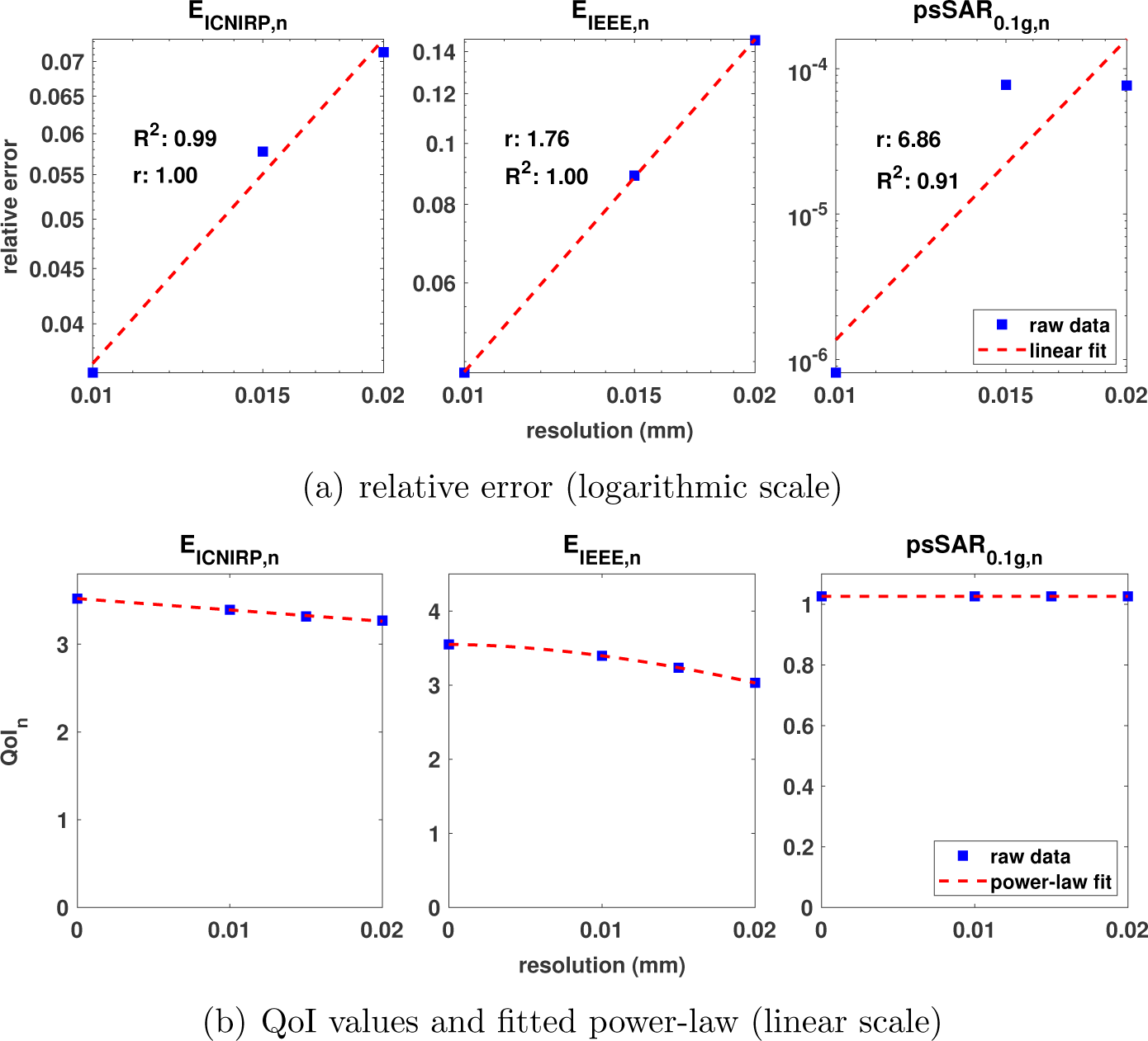
Convergence analysis for implant-perpendicular field incidence: Discretization convergence analysis for the dosimetric quantities of interest (QoIs) E_ICNIRP,n_, E_IEEE,n_, and psSAR_0.1g,n_. (a) Log-log plots of relative discretization errors as a function of resolution; the linear regression slope is indicative of the order of the numerical scheme. (b) Power-law fit to QoI, based on the linear regressions in the log-log plots. The interplay between these two views can be used to determine asymptotic QoI values at infinitely fine resolution (see Sec. 2.2.4.)

### 3.2. Dosimetric Enhancement Factors

*Generic implant* The QoIs (see Sec. 2.1) were computed for incidence angles in the range 0–90° degree in steps of 7.5°. The spatial distributions of E_IEEE,n_, psSAR_0.1g,n_, and ΔT_max,n_ are shown in Fig. 7 for a sample implant (*d*=0.8 mm, *L_c_*=3.0 mm, *L_i_*=0.5 mm), for parallel, 45° and perpendicular field incidence. The E_ICNIRP,n_ distribution is not shown, as it closely resembles that of E_IEEE,n_. The maximal E_ICNIRP,n_ and E_IEEE,n_ are found for *L_c_*=3.0 mm, *L_i_*=0.5 mm, and *θ*=15°, except for *d*=1.3 and 1.4 mm where E_IEEE,n_ is maximal at *θ*=7.5°. The results are summarized in Table 1 and visualized in Fig. 8(a) and Fig. 8(b). These figures show how E_IEEE,n_ varies as a function of incidence angles and contact or spacer length (the other parameters are fixed at the value which results in the maximum E_IEEE,n_ for the implant with 0.8 mm diameter (see Fig. A.2 for corresponding E_ICNIRP,n_ data). Figure. 9 compares E_ICNIRP,n_ and E_IEEE,n_ enhancement factors for different implant diameters. Figure. 8(c) and Fig. 8(d) show the same as Fig. 8(a) and Fig. 8(b), but for the psSAR_0.1g,n_ enhancement. It should be noted that the maximum of psSAR_0.1g,n_ is obtained at *θ* = 0°.

**Figure 7.**
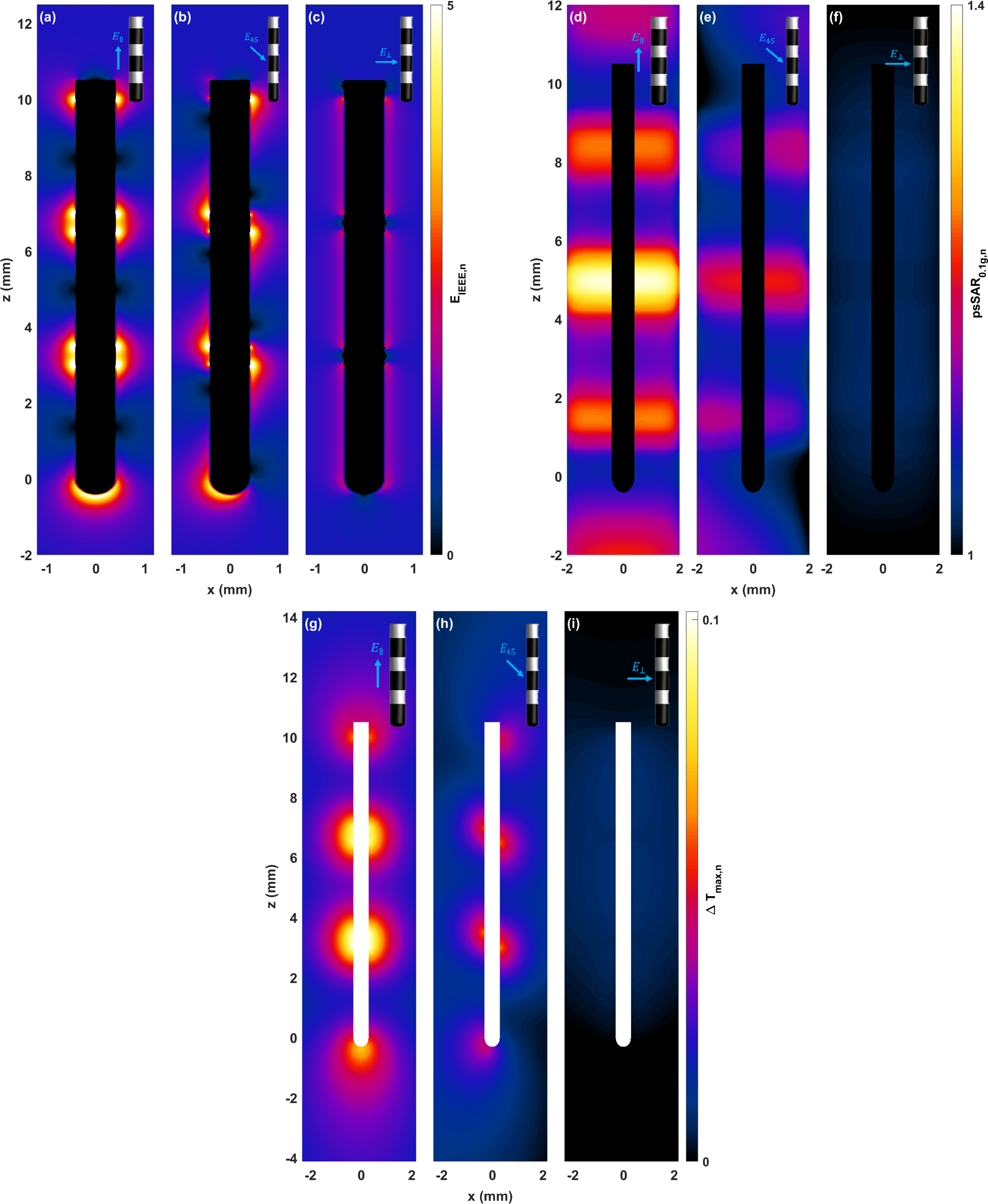
Distribution of quantities of interest for the generic implant with 0.8 mm diameter (*L_c_*=3.0 mm, *L_i_*=0.5 mm). E_IEEE,n_ distribution when the field is (a) parallel to the implant, (b) at a 45° incidence angle, and (c) perpendicular to the implant. psSAR_0.1g,n_ distribution when the field is (d) parallel to the implant, (e) at a 45° incidence angle, and (f) perpendicular to the implant. ΔT_max,n_ distribution when the field is (g) parallel to the implant, (h) at a 45° incidence angle, and (i) perpendicular to the implant.

**Figure 8.**
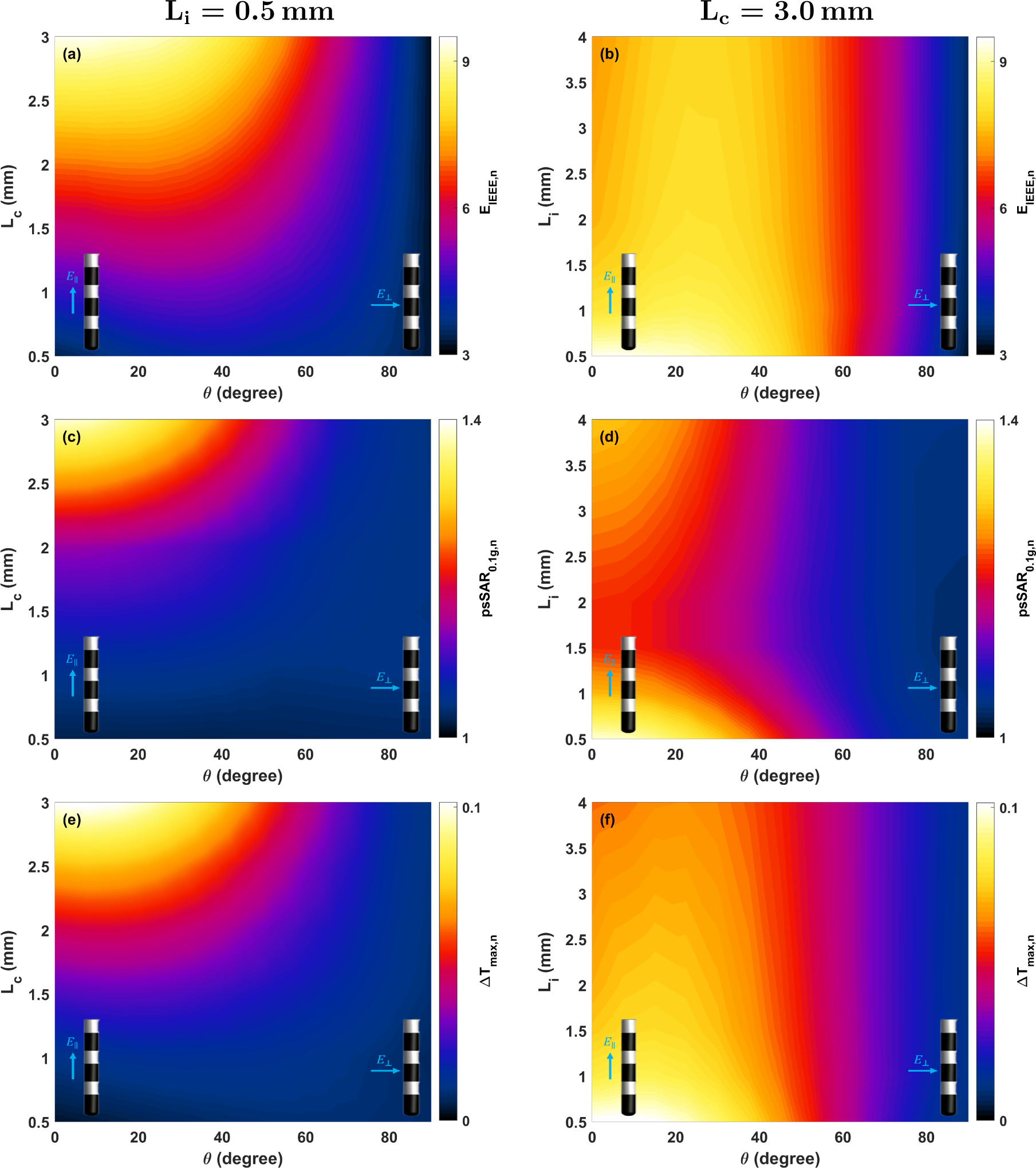
Quantitaties of interest illustration for the 0.8 mm diameter implant as a function of field-incidence angle, considering various contact and insulator lengths. Top: E_IEEE,n_ as a function of (a) contact length (fixed insulator length *L_i_*=0.5 mm) and (b) the insulator length (fixed contact length *L_c_*=3.0 mm). Middle: psSAR_0.1g,n_ as a function of (c) varying contact length (with fixed insulator length *L_i_*=0.5 mm) and (d) varying insulator length (with fixed contact length *L_c_*=3.0 mm). Bottom: Field-incidence angle dependence of ΔT_max,n_ as a function of (e) contact length (fixed insulator length *L_i_*=0.5 mm) and (f) the insulator length (fixed contact length *L_c_*=3.0 mm). For a comprehensive 3D visualization, please see Fig. A.3, Fig. A.4 and Fig. A.5.

**Figure 9.**
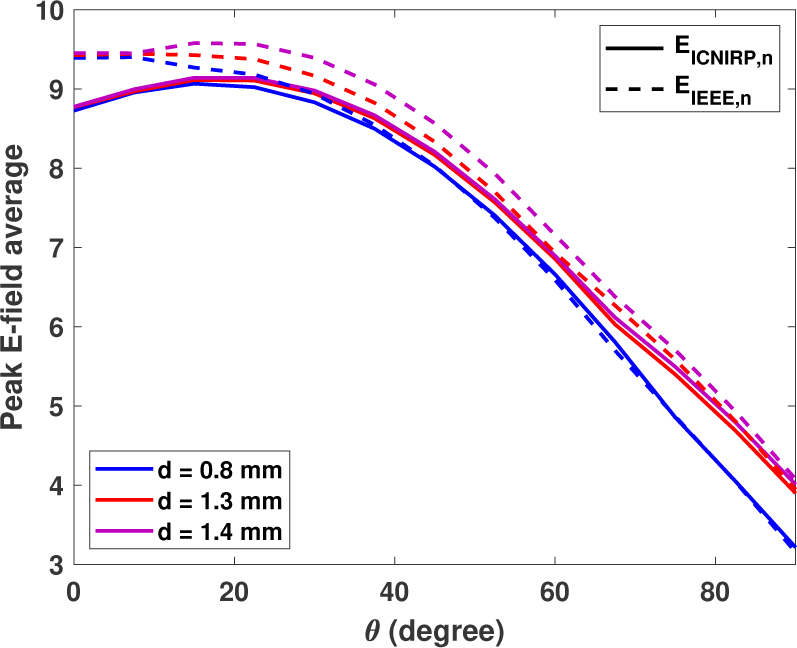
Impact of the E-field averaging scheme (solid lines: E_ICNIRP,n_, dashed lines: E_IEEE,n_) on the enhancement factors as a function of the field-incidence angle.

**Table 1.** Maximum E_ICNIRP,n_, E_IEEE,n_, psSAR_0.1g,n_ enhancement factors, and ΔT_max,n_ compared to the background for the studied implant geometries (d: diameter).

| d (mm) | $E_{\text{ICNIRP},n}$ | $E_{\text{IEEE},n}$ | $\text{psSAR}_{0.1g,n}$ | $\Delta T_{\text{max},n}$ |
| --- | --- | --- | --- | --- |
| 0.8 | 9.1 | 9.4 | 1.39 | 0.10 |
| 1.3 | 9.1 | 9.4 | 1.59 | 0.12 |
| 1.4 | 9.1 | 9.6 | 1.64 | 0.13 |

As Fig. 9 shows, E_IEEE,n_ provides more conservative enhancement factors compared to E_ICNIRP,n_. Therefore, we focus on it throughout the rest of the analysis; E_ICNIRP,n_ results are provided in Appendix A.

The results of varying the averaging length in E_IEEE,n_ and E_ICNIRP,n_ are presented in Table 2. As expected, increasing the averaging length reduces the E_ICNIRP,n_ and E_IEEE,n_ as a result of the highly localized nature of the field enhancement.

**Table 2.** Maximum of E_ICNIRP,n_ and E_IEEE,n_ enhancement factors for the studied implant geometries compared to the background as a function of averaging length (d: diameter).

| d (mm) | $E_{\text{ICNIRP},n}$ | | | $E_{\text{IEEE},n}$ | | |
| --- | --- | --- | --- | --- | --- | --- |
|  | 0.1 mm | 0.15 mm | 0.2 mm | 0.1 mm | 0.15 mm | 0.2 mm |
| 0.8 | 9.1 | 7.3 | 6.3 | 9.4 | 8.2 | 7.5 |
| 1.3 | 9.1 | 7.4 | 6.4 | 9.4 | 8.3 | 7.6 |
| 1.4 | 9.1 | 7.5 | 6.4 | 9.6 | 9.3 | 7.7 |

*Segmented and HD electrodes* For segmented and HD electrodes (see Fig. 3(e-h)), E_IEEE,n_ and E_ICNIRP,n_ were computed for *θ* = 0 *−* 90° and *ϕ* = 0 *−* 120° in steps of 7.5°. The results for E_IEEE,n_ are presented in Fig. 10(a-c) (refer to Fig. A.6 for E_ICNIRP,n_). As the results indicate, the maximum E_IEEE,n_ enhancement is found for the Boston Scientific Cartesia DBS lead model, where 7.0 can be reached – a value below the worst case of the generic implant geometry (see Table 1). However, when comparing the segmented electrodes with a similar implant lacking segmentation, they exhibit a higher enhancement. To explore this further, we segmented the middle contact of the worse case generic implant geometry (i. e. *d*=1.4 mm, *L_c_*=3.0 mm, and *L_i_*=0.5 mm). Under this condition, we observed that the maximum E_IEEE,n_ increases from 9.6 to 10.7, representing a 10 % increase, which might be related to the close proximity of PEC edges and an overlap of associated enhancements. The results are depicted in Fig. 10(d) (and Fig. A.6(d) for E_ICNIRP,n_) for different angles of incidence of E-field.

**Figure 10.**
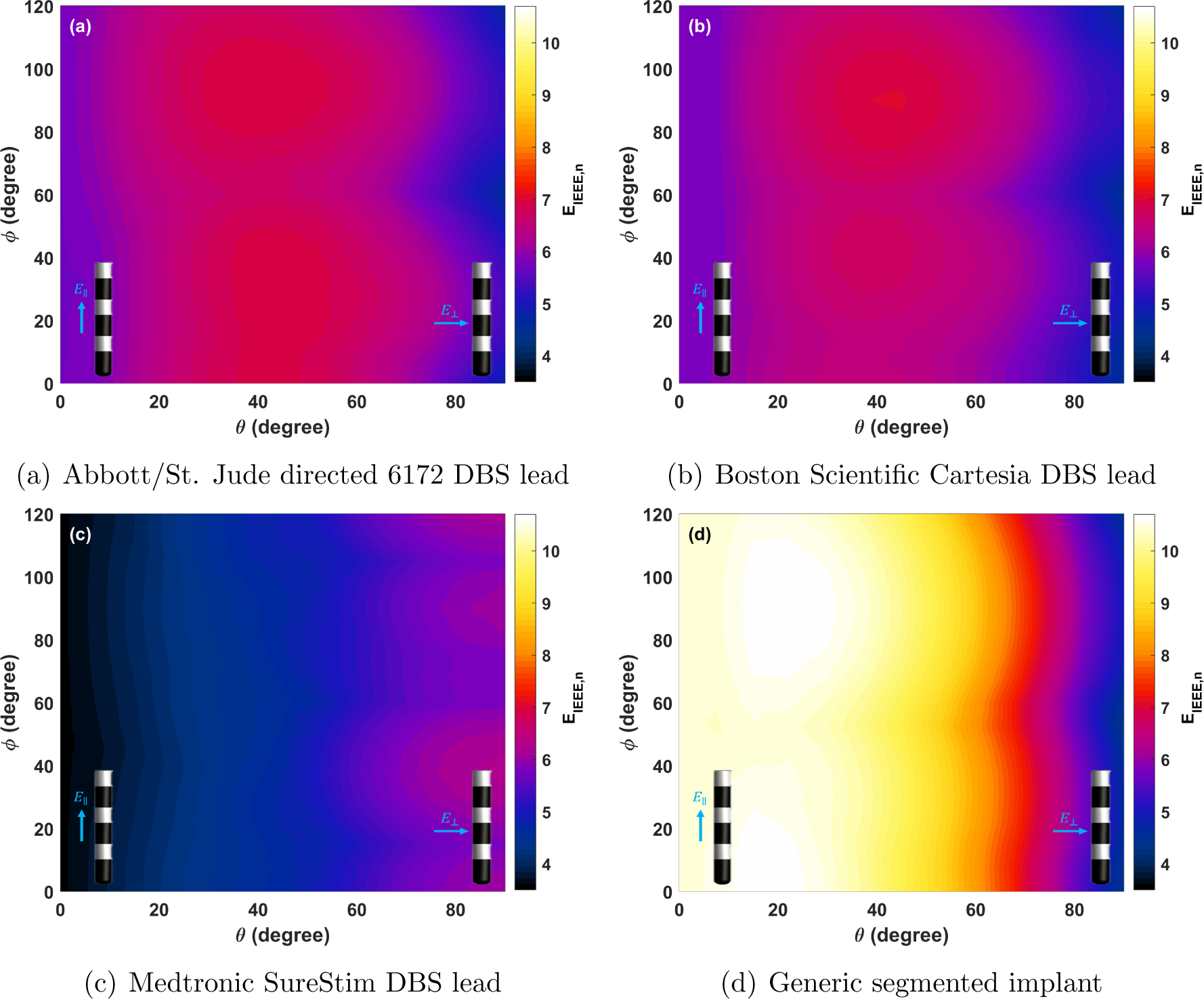
(a-d) E_IEEE,n_ for the segmented and HD electrode leads (see Fig.3(e-h)) as a function of field-incidence angles. The generic segmented implant (d) is based on the unsegmented-worst-case generic implants geometry (*d*=1.4 mm, *L_c_*=3.0 mm, and *L_i_*=0.5 mm).

*Elongated conductive cylinders* E_IEEE,n_ and E_ICNIRP,n_ were computed for elongated conductive cylinders (see Fig. 3(i))– both to study the impact of conductor length and to consider passive implants (e.g., cerebrovascular stents). Fig. 11(a) shows E_IEEE,n_ obtained when varying the cylinder diameter and length from 1 to 5 mm (in steps of 1 mm) and from 5 to 50 mm (in steps of 50 mm), respectively (see Fig. A.8(a) for corresponding E_ICNIRP,n_ results). The impact of length and diameter on E_IEEE,n_ is shown in more detail in Fig. 11(b) and Fig. 11(c) (see Fig. A.8(b) and Fig. A.8(c) for corresponding E_ICNIRP,n_ results). The maximum E_IEEE,n_ occurs when the E-field is almost parallel to the implant (between 0 and 22.5°, depending on implant’s length). E_IEEE,n_ increases linearly with implant length (see Fig. 11(b)) and decreases with implant diameter (approximately proportional to *d^−^*^0.4^; see Fig. 11(c)). For the detailed stent geometry (see Fig. 3(k)), an E_IEEE,n_ of 18 was found.

**Figure 11.**
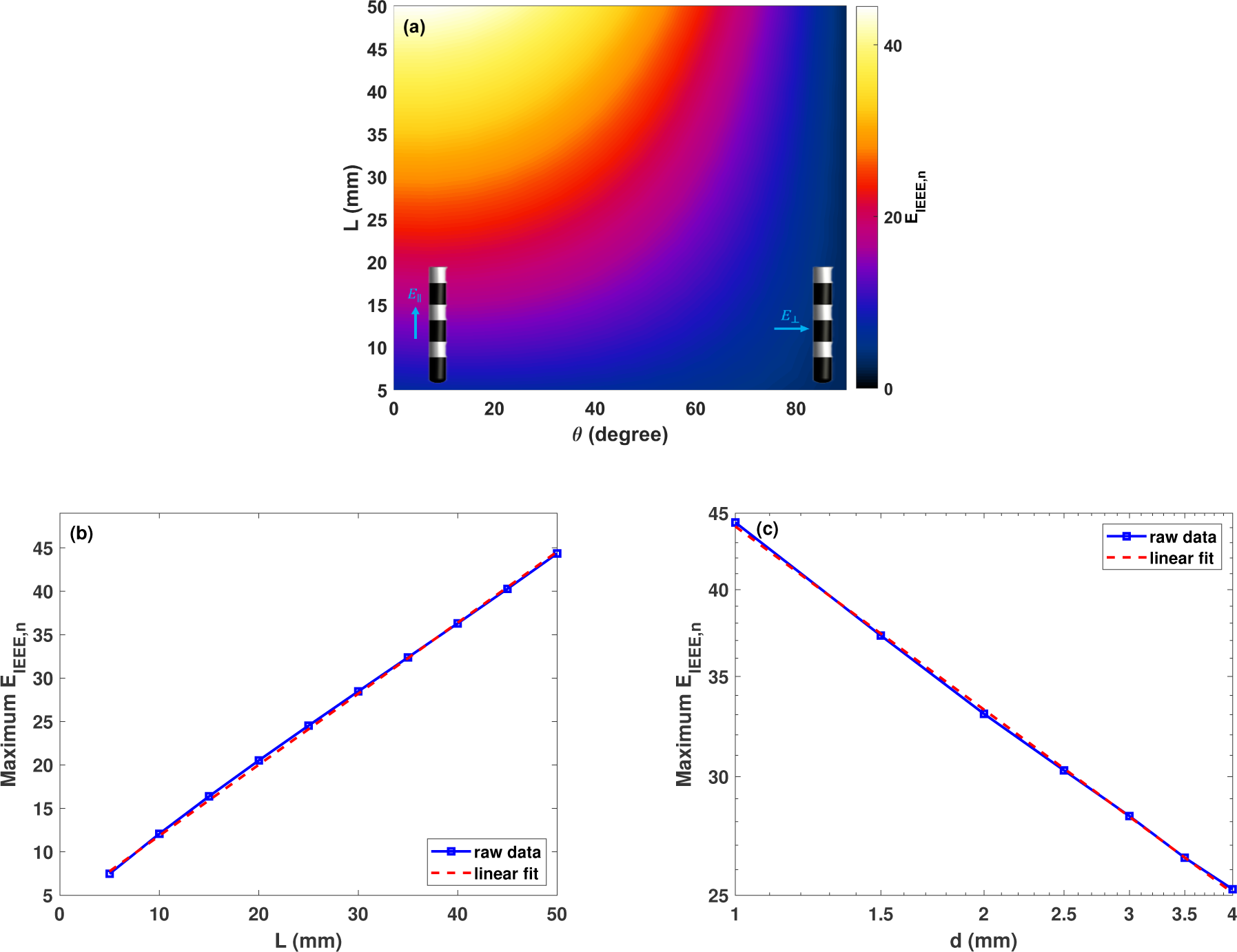
(a) E_IEEE,n_ for elongated cylindrical implants with 1 mm diameter as a function of their length and the field-incidence angle. The maximal E_IEEE,n_ for elongated cylindrical implants is illustrated in (b) as a function of length when the diameter is 1 mm (linear regression: maximal E_IEEE,n_ *≈* 0.8 *mm^−^*^1^*L* + 3.7) over all *θ*, and (c) as a function of the implant diameter when the length is 50 mm (power law fit: E_IEEE,n_ *≈* 44 *mm*^0.41^ *d^−^*^0.41^). Note the logarithmic *x* - and *y* -axis scales in (c).

*Scar tissue encapsulation* E_IEEE,n_ and E_ICNIRP,n_ were computed for generic implant geometries (as described in Sec. 2.2.1) with added scar tissue encapsulation (Fig. 3(j)). The scar tissue thickness and conductivity were varied from 25–300 *µ*m and assigned conductivites of 0.019 and 0.05 S/m. Figure. 12 illustrates E_IEEE,n_ for all scar tissue thicknesses as a function of the E-field angles of incidence (*θ*). These results are presented for the scarless-worst-case implant geometry (*L_c_*=3.0 mm, *L_i_*=0.5 mm), as derived in Sec. 3.2. The corresponding results for E_ICNIRP,n_) are shown in Fig. A.7. The maximum E_IEEE,n_ across all scar tissue thicknesses are shown as a function of E-field angle of incidence in Fig. 12(a) for different implant diameters and both conductivity values. The maximum brain tissue E_IEEE,n_ is reduced from 9.6 to 9.2 in the presence of 25*µ*m scar tissue. The maximum E_IEEE,n_ now occurs at *θ* = 45°, while in the scarless case (close to) axis-aligned field orientation produces the highest enhancement (see Sec. 4.1). It is important to note that the E-field within the scar tissue was excluded in this analysis, because no scar tissue stimulation is expected and scar tissue is far less temperature sensitive than brain tissue. If it was included, the maximum E_IEEE,n_ would increase from 9.6 to 24.2 (see Fig. A.7(c), Fig. A.7(d) and Fig. A.7(e)).

**Figure 12.**
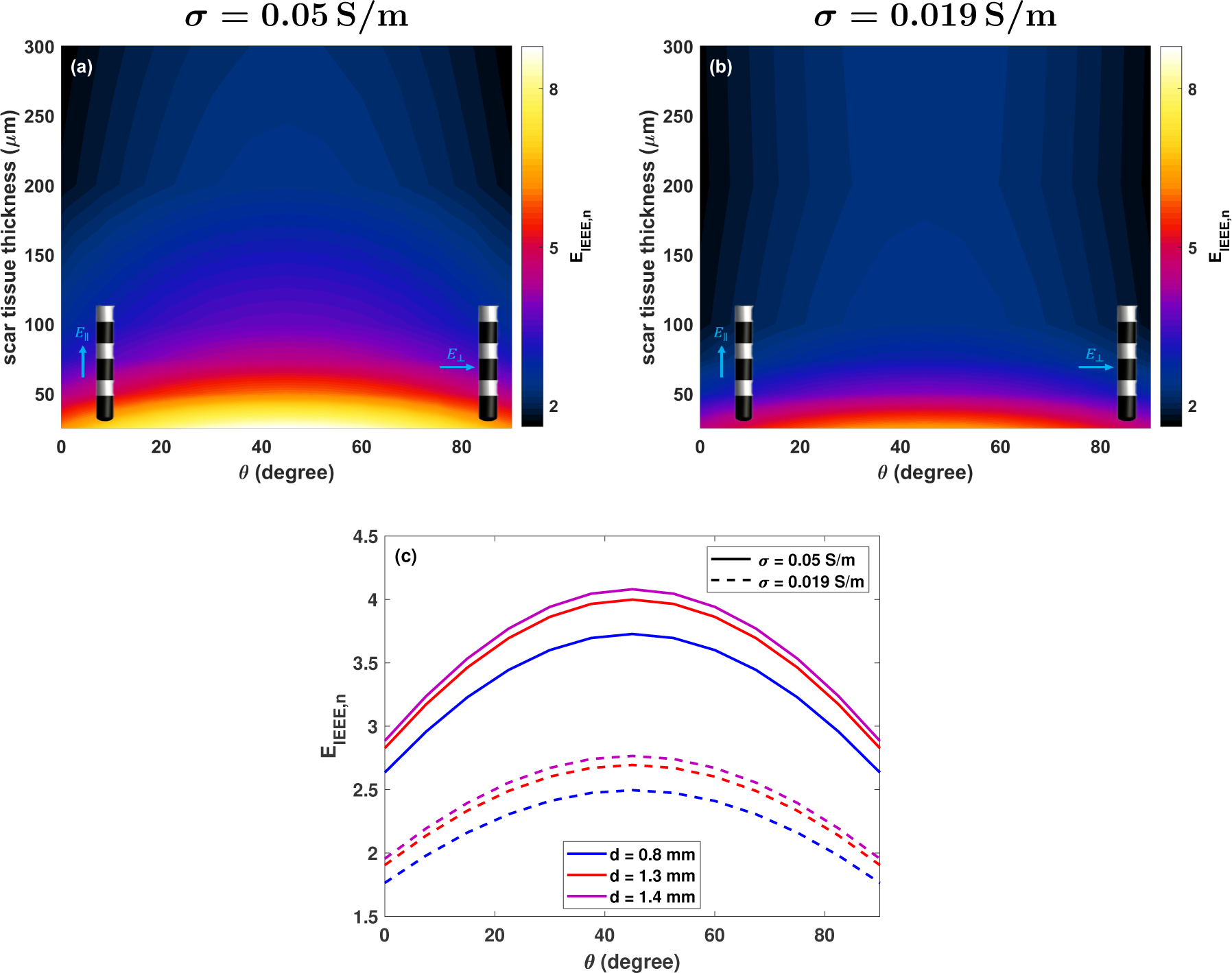
(a,b) E_IEEE,n_ as a function of field-incidence angles for the worst case generic implant geometry (0.8 mm diameter, *L_c_*=3.0 mm, *L_i_*=0.5,mm) encapsulated in scar tissue (Fig.3(j)). Note that the E-field in scar tissue is excluded from this analysis. (c) Maximum E_IEEE,n_ enhancement as a function of implant diameter for 100 *µ*m scar tissue thickness. Note that the E-field in scar tissue is excluded from this analysis.

### 3.3. Anatomical Model Validation of Enhancement Factor Approach

The maximum estimated E-field obtained from the simulation without SEEG implants, using the enhancement factors derived from the generic implant simulation, was plotted against the maximum E-field computed directly from the FEM simulation with integrated SEEG electrodes (see Fig. 13). Note that the comparison is not ideal, as the enhancement factors predict IEEE-averaged peak field, which could not be extracted from the full anatomical simulation, because the IEEE-averaging algorithm is not available for unstructured discretizations (tetrahedral). Instead, the peak E-fields from the FEM simulation are used. The results indicate a strong correlation (*R*^2^ =0.79 and 0.87 for the tDCS montages in Fig. 4(a) and Fig. 4(d), respectively) between the estimated and simulated values. The magenta line in the figure represents the ideal case. The estimated E-field tends to be higher than that obtained in full simulations. The maximal over and under-estimation factors are 2.3 and 0.6. The RMS relative difference between estimations and simulations are 60 % and 30 % for the tDCS montages in Fig. 4(a) and Fig. 4(d). The results using the enhancement factors obtained with 0.2 mm averaging was also presented in Fig. 13 to match the simulation discretization resolution. The maximal over and under-estimation factors are now 1.7 and 0.4. The RMS relative difference between estimations and simulations are 30% and 20% for the tDCS montages in Fig. 4(a) and Fig. 4(d).

**Figure 13.**
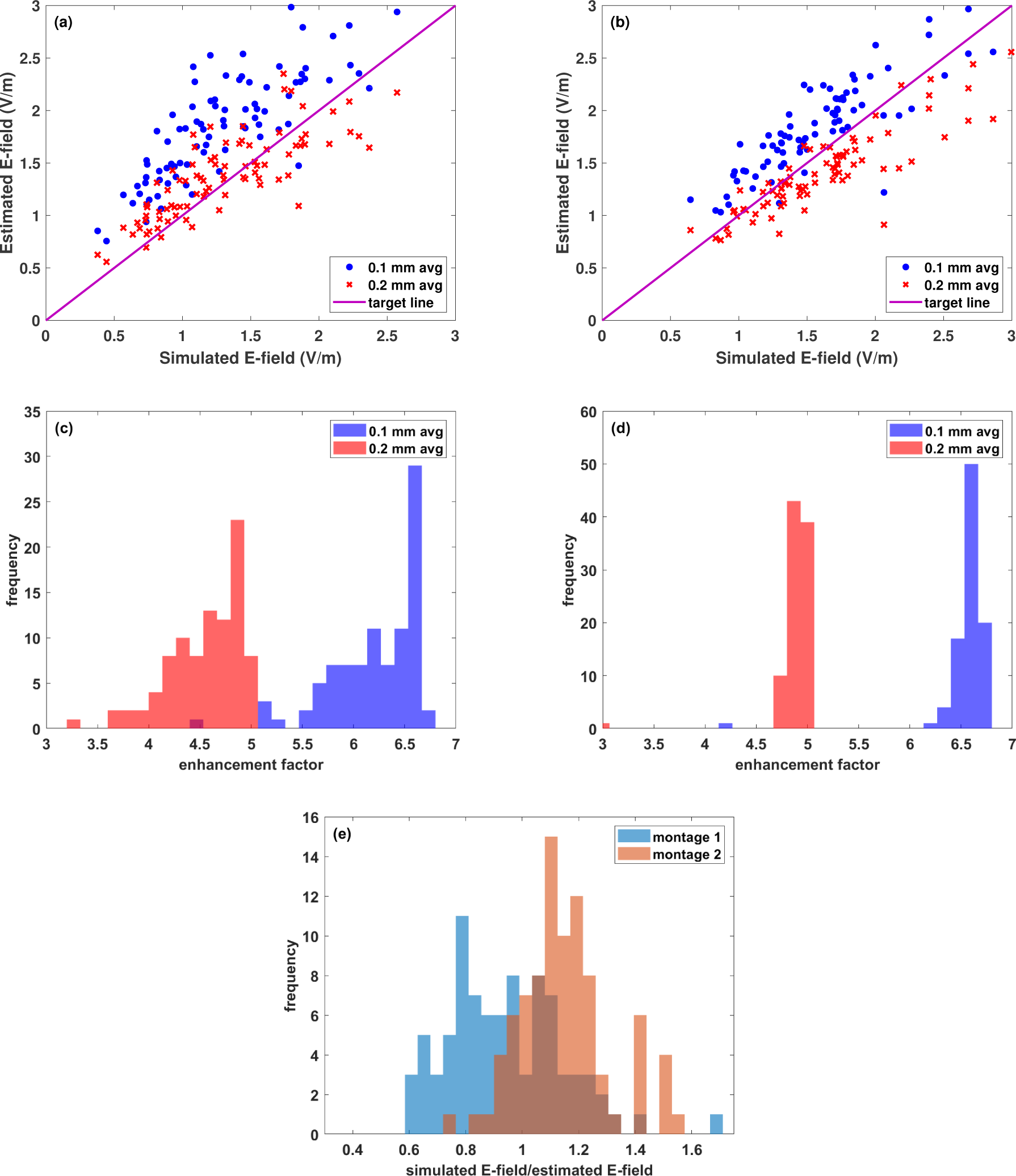
(a,b) Predicted and simulated E-field peak magnitudes near SEEG electrode contacts for the tDCS montages presented in Fig. 4(a) and Fig. 4(d), respectively. The full simulations featured a detailed anatomical head model with implanted SEEG leads (see Fig. 4), while the predictions were based on simulations without implants and the direction-dependent enhancement factors obtained using the simplified setup (the histogram of the enhancement factors are depicted in subfigures (c) and (d) of Fig. 4(a) and Fig. 4(d), respectively). The magenta line in the figure represents ideal agreement. (e) Histogram of the ratio between simulated and estimated E-field using 0.2 mm average length.

### 3.4. Tissue Heating

Steady-state temperature increase simulations were performed for all generic implant geometries (see Sec. 2.2.1) and the incident E-field dependence was assessed as described in Sec. 2.3. The results are presented in Fig. 8(e) and Fig. 8(f) for a 0.8 mm diameter implant as a function of E-field angle of incidence and contact or spacer length (other parameters are fixed at the value which results in the maximum temperature rise). Fig. 14(a) provides a comparison of temperature increases for implants with different diameters across varying E-field angle of incidence. As the results show, ΔT_max,n_ always remains below 0.11. Furthermore, it is important to highlight the strong correlation between ΔT_max,n_ and psSAR_0.1g,n_ evident in Fig. 14(b) (0.8 mm diameter; worst case implant scenario: *L_c_*=3.0 mm, *L_i_*=0.5 mm). The linear correlation coefficient between ΔT_max,n_ and psSAR_0.1g,n_ was computed for each combination of *L_c_* and *L_i_*, resulting in 0.69*±*0.19 for implants with 0.8 mm diameter.

**Figure 14.**
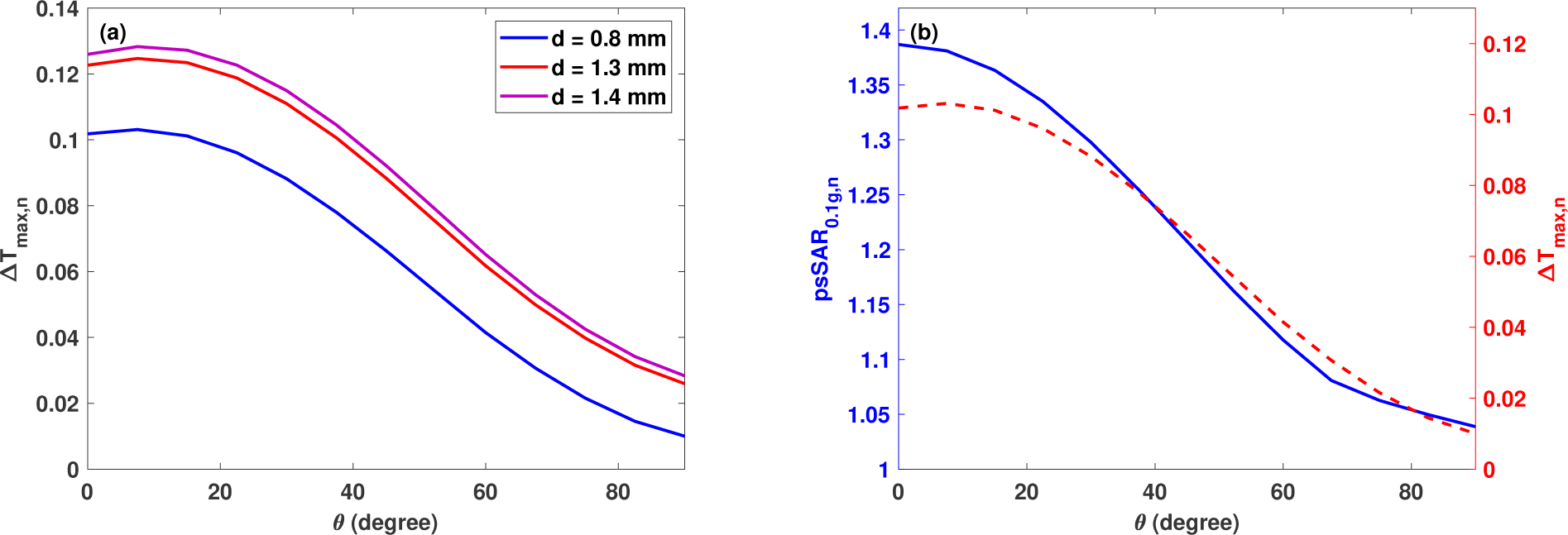
(a) ΔT_max,n_ for the worst case generic implant (*L_c_*=3.0 mm, *L_i_*=0.5 mm) as a function of the field-incidence angle. (b) ΔT_max,n_ and psSAR_0.1g,n_ for the worst case generic implant (*L_c_*=3.0 mm, *L_i_*=0.5 mm) with 0.8 mm diameter as a function of the field-incidence angle.

**Figure 15.**
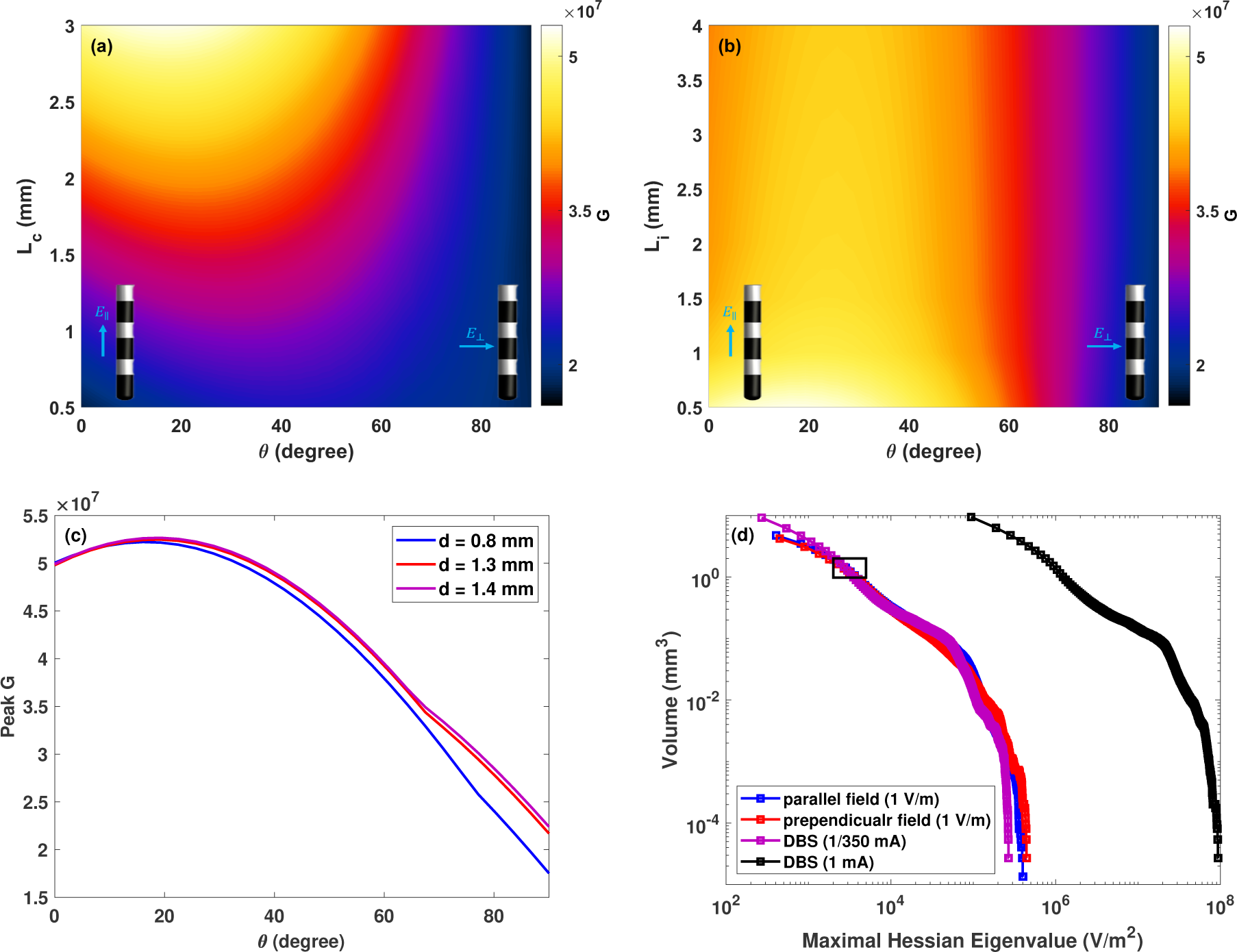
The peak Hessian Eigenvalue (absolute value) for a 0.8 mm diameter implant as a function of (a) contact length (fixed insulator length *L_i_*=0.5 mm) and (b) the insulator length (fixed contact length *L_c_*=3.0 mm). (c) *G* as a function of the field-incidence angle for the worst case generic implant (*L_c_*=3.0 mm, *L_i_*=0.5 mm). (d) Volume of tissue exposed to at least a certain field gradient magnitude (can be interpreted in terms of volume of tissue activated) for the 0.8 mm diameter implant (*L_c_*=0.5 mm and *L_i_*=0.5 mm) in passive mode (subjected to 350 V/m radially and tangentially oriented incident E-fields) and active mode (1 mA input current applied in monopolar cathodic configuration). The E-field rescaling to 350 V/m ensures that the field gradient strengths isolevel encompassing 1 *mm*^3^ is identical, enabling qualitative comparison of the neuromodulatory impact of passive field enhancement to active DBS stimulation.

### 3.5. Impedance Measurements

Consider first the macro-micro depth electrode lead with 6 macro (stimulation) electrodes and 10 micro monitoring electrodes. The capacitance between electrodes is in the range 18 – 20 pF or 16 – 17 pF for macro and micro electrodes respectively. Because the lead splits after 25 cm the capacitance between the leads of the micro and macro conductors is less at approximately 12 pF. The material properties of the surrounding tissue minimally affect the overall capacitance between electrodes as the measured value is the result of all inter conductor capacitances. The lead conductor capacitance to the tissue, for 15 cm length of lead embedded in conductive medium is approximately 14 pF. For the Medtronic SenSight directional lead, the capacitance is generally in the range 50 – 80 pF. The capacitance to the tissue is approximately 44 pF for any connection. The impedance of the leads alone is one element and unless they are disconnected from any device it cannot be considered in isolation. A lead connected to a device – whether recording or stimulating – will have additional current pathways. In the case of recording devices, these pathways are likely to be high impedance, unless MRI or electromagnetic interference (EMI) filtering has been included. In the case of stimulating devices, the impedance depends on the employed architecture in addition to any MRI or EMI filtering. MRI or EMI filtering typically introduces additional capacitance in the range of nF to tens of nF between contacts, resulting in a much higher capacitance. The impedance of the ohmic pathway through brain tissue for the two aforementioned leads was determined through simulations, yielding approximately 500,Ω between adjacent contacts for the macro-micro depth electrode lead and 900–1370,Ω for the Medtronic SenSight directional lead (range reflects differences between segmented and ring contacts).

### 3.6. Neurostimulation

#### 3.6.1. Worst Case Gradient

The *G* was computed for all generic implants and the results for the 0.8 mm diameter implant are shown in Fig. 15(a) and Fig. 15(b). For all implant diameters, the highest *G* is found for *L_c_*=3.0 mm, *L_i_*=0.5 mm, and *θ*=15°. Fig. 15(c) compares the peak *G* between different implant diameters.

#### 3.6.2. Comparison with DBS Activation

The stimulation ability of passively exposed implants due to localized E-field enhancement was quantitative compared to that of DBS-like active implants in monopolar cathodic configuration. As i) the high local enhancements near electrode edges are similarly shaped and ii) the cumulative dose-volume histograms (DVH) of the maximal field gradient distributions (*G*) are nearly identical up to a constant scale factor (see Fig. 16 and Fig. 17), it is reasonable to interpret the results in terms of a relative scaling factor that renders the two configurations equivalent in terms of volume of tissue activation (VTA; a common metric in DBS impact analysis). Fig. 15(d) compares the field gradient

**Figure 16.**
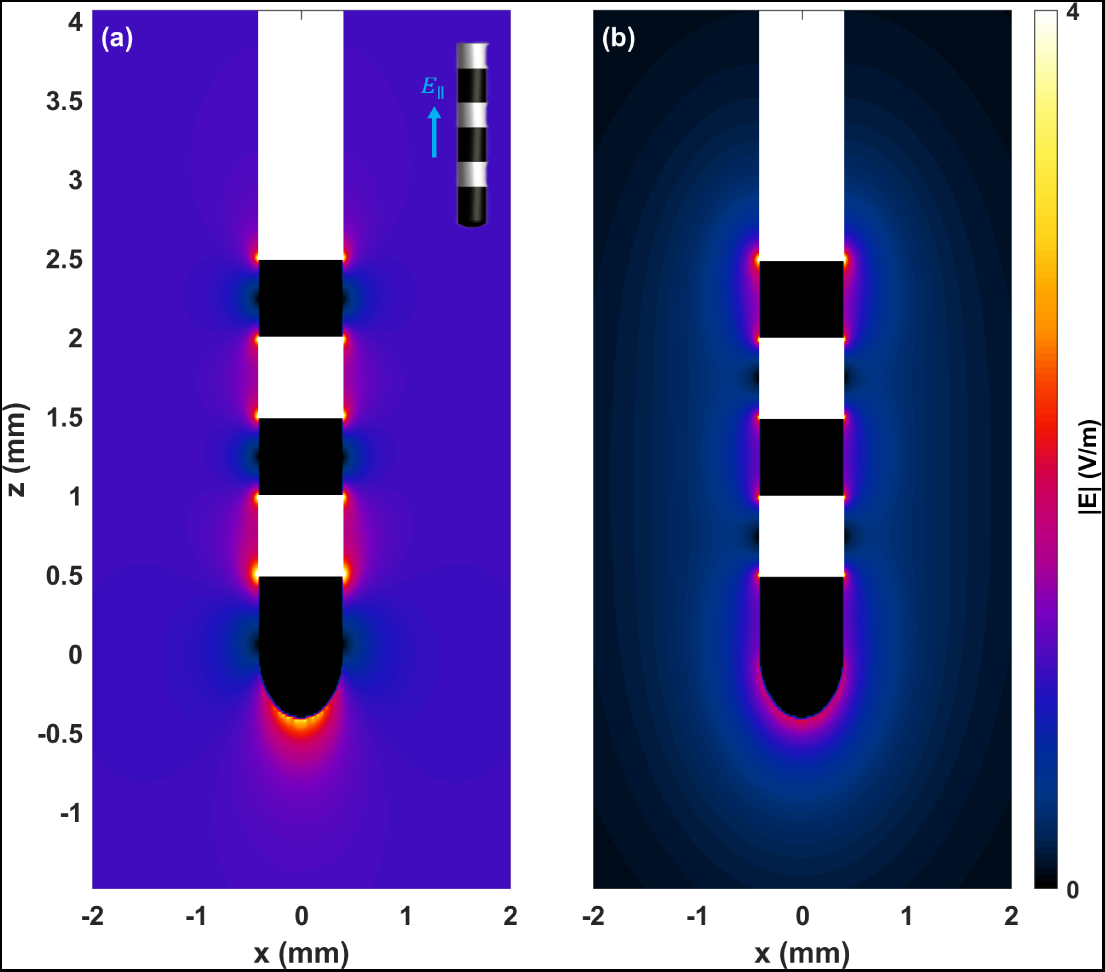
Comparison between the E-field distribution for (a) the generic implant (*d*=0.8 mm, *L_c_*=0.5 mm, *L_i_*=0.5 mm) exposed to an incident E-field of 1 V/m parallel to the implant axis and (b) the same implant in monopolar stimulation configuration with an input current of 1/350 mA - the return electrode is assumed to be very far away.

**Figure 17.**
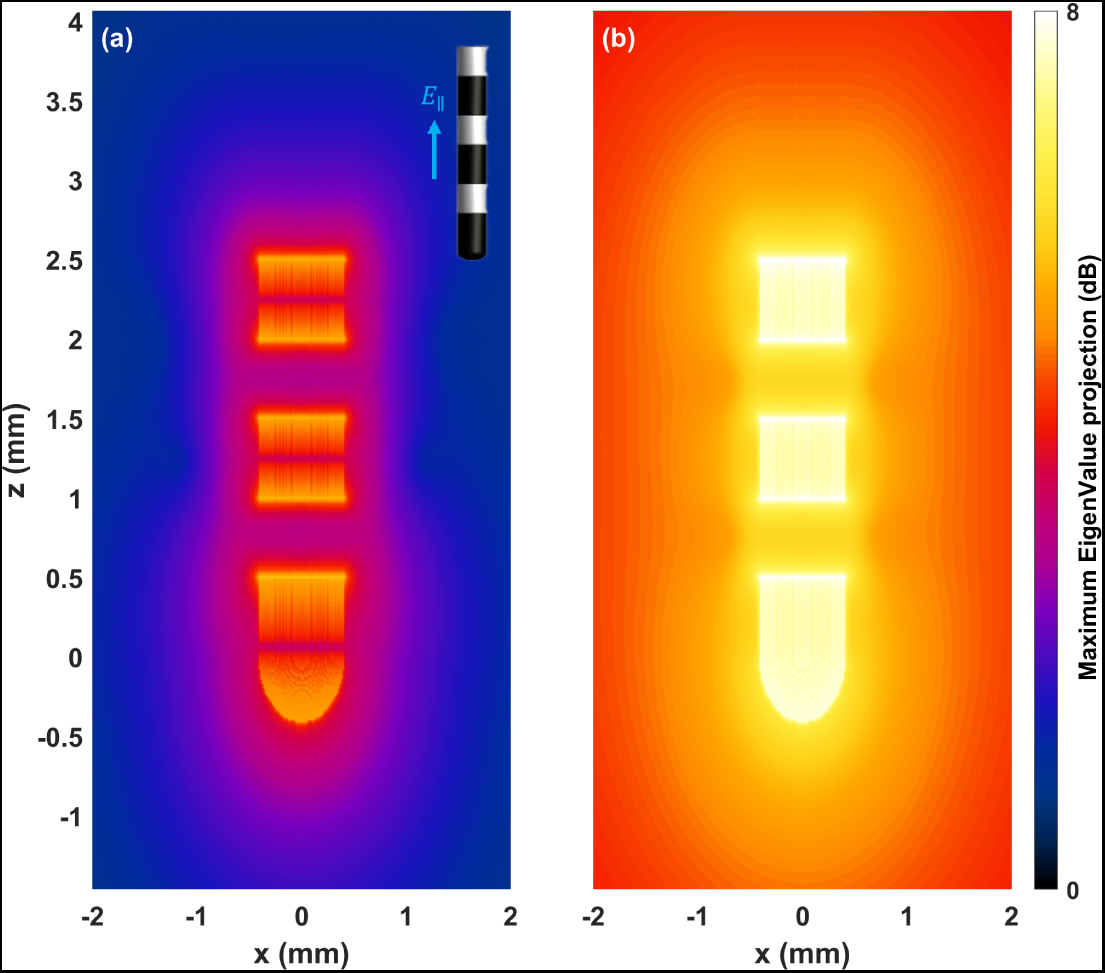
Comparison between the maximum Eigen value projection distribution for (a) the generic implant (*d*=0.8 mm, *L_c_*=0.5 mm, *L_i_*=0.5 mm) exposed to an incident E-field of 1 V/m parallel to the implant axis and (b) the same implant in monopolar stimulation configuration with an input current of 1/350 mA - the return electrode is assumed to be very far away.

DVHs of the actively driven implant (1 mA input current) and the enhancement resulting from passive exposure to a radially oriented 1 V/m incident E-field. It also shows the passive DVH rescaled such that the 1 *mm*^3^ VTA gradient strength is equal to that of the active configuration. The comparison suggests similar stimulation ability of a 1 mA input current DBS and the enhancement resulting for the same electrode at an incident E-field magnitude of about 350 V/m.

### 3.7. Abandoned Lead Simulation

Figure. 5(e,f) show the E-field distribution from the abandoned lead setup in Sec. 2.6. The total current exiting through the implant contacts is 20 % (respectively 40 %) of the applied tES current for the 0.2 mm (respectively 1 mm) wire. This use-case also represents leads with damaged insulation.

## 4. Discussion

### 4.1. EM and Thermal Exposure Enhancement

*Local field enhancement* The results reported in Sec. 3.2 are in accordance with expectations from a simple mechanistic picture. Metallic contacts act as low impedance pathways. Currents are diverted locally to enter at the closest end, short-circuit through the metallic contact, and exit at the opposite end, thereby increasing the current density at both ends. If the incident field orientation is not parallel to the implant axis, a polarization ‘quadrupole’ results. Field-line-orthogonality to PEC and related edge effects further contribute to locally enhancing the fields at these poles. This effect increases with the extent of the PEC along the field exposure direction. Therefore, enhancement factors increase nearly linearly with conductor length (most evident in the data from the generic passive implant; see Fig. 11(b)). The worst-case is therefore observed for the longest contact length at an orientation where the contact diagonal is almost oriented with the field (see Fig. 8 and Fig. 9). Usually, the contact diagonal is close to axial and, consequently, maximal enhancement occurs for near-parallel incidence – however, this is not the case for the shortest contact length, where the diagonal orientation is closer to radial and the maximal enhancement is observed for larger incidence angles. For most electrode separation distances, the contacts act as independent enhancers. Only for the closest separations (around and below 1 mm) is an additional enhancement apparent (see. Fig. 8(b), Fig.8(d), and Fig. 8(f)), which can be due to overlap of the field enhancements from adjacent edges, or to coupling between the contacts that produces an effectively longer conductor. The diameter (circumference) dependence is weaker – fitting a power-law to the data from the elongated passive implant suggests an exponent of *−*0.41. This can be interpreted as an interplay between current distributing over a larger circumference (reducing the current density and thus E-field) and the resistance to current entering and leaving the PEC simultaneously decreasing (thus increasing the current density and E-field). As these two effects partly compensate, a weak diameter dependence results.

*Peak spatial SAR enhancement* psSAR_0.1g,n_ shows a similar behavior (see Fig. 8(c) and Fig. 8(d)), but are much smaller (up to 1.4, compared to the factor 100 that could have been expected from naively squaring the E-field enhancements). This is likely due to the increased averaging volume, which dilutes the highly localized enhancement. The psSAR_0.1g,n_ behavior closely corresponds to that of the temperature increase (see Fig. 14(b) and Sec. 3.4), supporting the choice of the SAR averaging mass.

*Temperature increase* Temperature increase only becomes tES-limiting with increasing frequency [4]. While the local field enhancement can increase the psSAR_0.1g,n_ by up to 40 %, the maximal temperature increase remains below 11 % of the background temperature increase – likely because of effective heat diffusion due to the high power-deposition gradients.

This implies that the additional brain temperature rise during tES stimulation in the presence of the implant is not a important additional concern.

*Scar tissue* Scar tissue encapsulation renders the parameter dependence more complex. This is due to the distinct behaviors of the parallel and perpendicular (relative to the scar-tissue interface) field components: Perpendicular current density (**J**) is preserved, but the corresponding E-field is discontinuous as a result of dielectric contrast. Conversely, parallel E-field components are continuous, while the associated **J** jumps. Their relative weight changes as a function of the incident angle, thus modifying the implant-geometry-dependent enhancement behavior. However, brain field enhancement in the presence of scar tissue is reduced compared to the scar-less case, such that the latter constitutes the relevant worst-case condition. When also considering the scar tissue exposure itself, the enhancement is higher, but considering the expected reduction in sensitivity and excitability, brain tissue exposure is expected to be limiting. In conclusion, while scar tissue formation around chronic implants can lead to further field magnitude increases within that tissue, the surrounding brain tissue exposure actually decreases.

*Anisotropy* To asses the impact of local anisotropy, additional simulations with tensorial conductivities were performed for the worst case scenario involving the generic implant with 0.8 mm diameter (*L_c_*=3.0 mm and *L_i_*=0.5,mm). The principal tensor axis was assigned the white matter axial conductivity value from the IT’IS database [56], while the radial value was assigned to the two perpendicular directions. That principal axis was once aligned with the implant and once perpendicular to it. Subsequently, the QoIs were extracted (see Fig. A.9 for E_IEEE,n_ results). The results indicate that the maximum E_IEEE,n_ increases from 9.4 to 10.0 with anisotropic conductivity, representing only a 6 % increase.

*Elongated passive implants* Longer conductive implants, such as stents, can become problematic at even lower exposure intensities, as E-field enhancements (and consequently *G*) scale linearly with conductor length (see Fig. 11(b)) and SAR and temperature increase scale quadratically. For the detailed stent geometry (see Fig. 3(k)), an E_IEEE,n_ of 18 was found, which is higher than the value of 10 obtained using the fitted diameter-and length-dependent enhancement estimation from Fig. 11. The latter is based on the data of the simplified elongated cylinder implant. In view of the 0.2 mm stent wire thickness, which produces more localized enhancement, the estimation was also performed using said wire diameter. In that case, an E_IEEE,n_ of 30 is estimated, such that the simulated value is between the predictions obtained using the stent and the wire diameters. Using the finer diameter is expected to always produce conservative estimations. For a more detailed exposure safety assessment approach for elongated passive implants, see the methodology of [60].

*Segmented & HD contacts* For some geometry parameters in the studied range, segmented electrodes are found to produce slightly higher enhancements than unsegmented ones. This is likely explained by edge proximity sharp corner effects. The HD contacts – as a result of their small dimensions – produce weaker enhancements than the cylindrical ones and do not constitute worst-case.

*Enhancement factor approximation* The acceptability of computing enhancement factors, thus separating incidence field computation from the computational of the local field distribution, was investigated. For that, simulations of a head model with integrated electrodes were compared to predictions obtained using a simulation without integrated electrodes in combination with the enhancement factors from the simplified setups. As previously mentioned (Sec. 3.3), the comparison is not ideal, as IEEE-averaged peak fields could not be extracted from the full anatomical simulation. Instead the peak E-fields from the FEM simulation are used. While the actual peak values are expected to be larger than the averaged quantities, the coarseness of the practically achievable mesh resolution reduces the field magnitude. When comparing the integrated simulation with a resolution of 0.2 mm to the 0.2 mm-averaged predictions, the RMS relative differences are 30% and 20% for the two tested montages (see Fig. 4). This supports the selected problem separation approach, provided a corresponding safety margin is included.

### 4.2. Capacitive Coupling

*Capacitive coupling into the lead* For a stimulation frequency of 100 Hz and a tES voltage of 1 V (current of 2 mA and impedance of 500 Ω) a worst-case current of 2.2 *µ*A (for the macro-micro depth electrode lead), respectively 6.9 *µ*A (for the Medtronic SenSight directional lead), is estimated using (5). This is 3 orders of magnitude weaker than a typical DBS current making the neurostimulatory effect comparable to that of the local field enhancement (see Sec. 3.6.2).

*Capacitive coupling between contacts* At a stimulation frequency of 100 Hz, the inter-contact impedances resulting from capacitive coupling between lead wires are in the order of 20 *M* Ω for the highest measured capacitance (i.e., 80 pF), which is more than four orders of magnitude larger than the impedance related to ohmic currents through tissue (around 500,Ω for the macro-micro depth electrode lead and 1240–1370,Ω for the Medtronic SenSight directional lead). Even at the highest stimulation frequencies of interest (100 kHz in TIS), the capacitive inter-contact impedances are still one order of magnitude above the ohmic ones. Therefore, capacitive coupling between contacts does not need to be considered with regard to safety or its impact on the current flow for leads similar to those measured in this study.

*Generality* Obviously, the capacitances are highly lead-design-specific and they should be remeasured for any specific implant of interest. Such measurements are simple to perform.

*Frequency dependence* Both forms of capacitive coupling result in effects that scale linearly with frequency. As a result, they become more relevant when the exposure frequency increases (see Sec. 4.4).

### 4.3. Ohmic Coupling (Abandoned or Damaged Leads)

In the investigated 0.2 mm wire scenario, 20 % of the applied tES currents enters the brain through implant contacts. Based on the equation 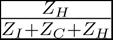 from Sec. 2.6, and the voltage level of the wire (close to the average of the tES electrodes), the resistance for entering the cut wire is in the order of 1-2 kΩ and similar in magnitude to the impedance between the implant contacts and the return electrode (pathway crosses the skull). The ratio of current entering the wire increases further for lower *Z_C_* (e.g., to above 40 % for the 1 mm wire). The magnitude of current flowing through the lead can result in DBS-like exposure conditions, resulting in a different stimulation paradigm than the intended tES. In consequence, tES should not be applied in the presence of abandoned or leads with damaged insulation.

### 4.4. Temporal Interference Stimulation

TIS deserves particular attention and care because:

the use of higher carrier frequencies means that higher currents are tolerated without triggering scalp sensations, typically resulting in the application of increased field strengths at depth;
while the increasing field strength might not necessarily be associated with an increased risk of unwanted neurostimulation, as stimulation thresholds also increase with frequency, the associated quadratic increase in energy deposition can lead to brain heating becoming the limiting safety concern [4];
furthermore, the stimulation-effectiveness of the TI LF modulation envelope is still unclear, such that an increased E-field magnitude might still be associated with safety-related neuromodulation concerns;
metallic conductors produce perfectly aligned (i.e, perpendicular) fields from the different channels, maximizing the risk of localized TI stimulation near implants (TI is expected to be most effective when the fields are aligned [9]);
with increasing frequency, capacitive coupling becomes proportionally more important, necessitating consideration of both capacitive current injection into the lead and reduced impedance pathways between electrode contacts; the former effect can reach problematic magnitudes (see Sec. 5.1).

### 4.5. TMS and Electroconvulsive Therapy

The present study specifically addresses tES exposure in the presence of implants. For *TMS* exposure, the *in vivo* exposure increases considerably. The local exposure enhancement factors are expected to remain similar (though additional validation is required). This, combined with the increased exposure, results in a corresponding increase in local field, SAR, and temperature increase near the metallic contacts (the latter two even quadratically with the field exposure). The mechanism of coupling into the leads also needs to be reconsidered, as inductive effects might become relevant.

*Electroconvulsive therapy (ECT)* is a psychiatric treatment where a generalized seizure is induced to manage refractory mental disorders [61]. It differs from tES in intensity, duration, and intended effects. While tES currents are typically in a range of 1-2 mA and last from several minutes to hour, ECT employs currents up to 800 mA in short bursts. Various studies [62, 63, 64, 65, 66] have investigated ECT safety in the presence of metallic objects, such as cerebral coils, deep brain stimulators, small metallic foreign body (e.g. bullets), cerebral clipping systems, metallic skull implants, and DBS implants. Based on over 40 reported cases in which no complications were observed, these studies conclude that with appropriate caution and safety precautions, psychiatrists may use it on an individualized basis. However, it should be noted that i) ECT is used to treat severe conditions, which warrants taking higher risks, ii) ECT exposes large parts of the brain and aims to trigger seizures, such that eliciting of severe neural responses is in fact desirable, and iii) while the quadratic increase in energy deposition as a function of current magnitude results in significantly increased heating rates, the application of ECT in short bursts contributes to significantly lowering the temperature increase, iv) the considered case studies are highly diverse and adverse effects would not necessarily have been manifested in evident behavioral changes (especially when considering the underlying severe disease conditions).

### 4.6. Assumptions and Limitations

The following assumptions and limitations apply to the present study:

*Considered implant geometries* The assessment of field enhancement utilized generic implant geometries, which are expected to comprehensively cover relevant SEEG and DBS geometries. However, it might still be desirable to reexamine specific implants when relevant. This particularly concerns passive implants, which were only coarsely modelled (in what is expected to mostly be a conservative approximation). The measurements performed to assess field coupling into the lead and low impedance pathways between contacts were obviously specific to the two tested implants and should be repeated for any implant of interest. Such measurements are straightforward to perform.

*Quasi-static approximation* The EM simulations were performed using an ohmic-current-dominated EQS solver. It employs approximations derived from the negligibility of displacement currents compared to ohmic currents and the small domain size in relation to the wavelength. The simulations did also not account for the frequency-dependence of dielectric properties (dispersion relationships). However, at the low frequencies of relevance to tES, these approximations are well justified, as confirmed by [67].

*Electrode-tissue-interfaces* Electrodes were modeled as being in direct ohmic contact with the tissue, neglecting capacitive and/or polarization effects. However, the employed simplification is expected to constitute a conservative case.

*Imperfect conductors* It has been suggested that conductors embedded within dielectric tissues and exposed to an E-field may exhibit a combination of perfect conductor and perfect insulator behavior, depending on the intensity of the applied field [68]. Simple circuit simulations indicate that this effect is not relevant for electrodes in (even high impedance) connection with measurement or stimulation electronics. However, for unconnected passive implants, like stents, the current investigations might overestimate field enhancement effects.

*Tissue heterogeneity and anisotropy* Tissues were modelled as homogeneous and in the anatomical head model case as isotropic. However, brain tissue can be anisotropic (especially white matter) and inhomogeneous. These factors can affect the E-field distribution (e.g., penetration depth) and the resulting neuromodulatory impact. However, while their impact on the incident field conditions can be high, the enhancement factors are not expected to change significantly for DBS/SEEG-like implants. However, the dielectric environment heterogeneity of implants situated in brain vasculature, which encompasses highly conductive blood, as well as epithelial vascular tissue, in addition to brain tissue, could have a major influence that must still be investigated.

*Dosimetric quantities-of-interest* The chosen dosimetric QoIs might be insufficient or suboptimal. For instance, the selected averaging length of 0.1 mm was chosen as a compromise between the distance between Ranvier nodes (a crucial factor in neurostimulation) and conservativeness in high field gradient locations. The averaging volume of 0.1 g was chosen to maximize correlations with tissue heating (larger averaging masses are used in the IEEE 1528 standard [36], for highly localized energy deposition, smaller averaging volumes or areas are needed; [69]). However, these choices might not produce the best correlations with safety-relevant tissue responses. Additionally, dendritic stimulation and cell polarization might be better predicted by directional field components.

We selected an averaging length of 0.1 mm. This choice is driven by a desire for conservatism in our approach and aligns with the length between two nodes of Ranvier in axon models, a crucial factor in neurostimulation.

*Multiple implants* Multiple implants in close proximity necessitate additional considerations, including potential further field enhancement resulting from overlapping local field enhancements or cross-coupling. Where relevant, such conditions should also be studied.

*Experimental confirmation* The present study relies on established forms of computational modeling that have been validated in related contexts. However, specific experimental validation of the here presented results, e.g., in measurement phantoms or *ex vivo* setups, while challenging, would be desirable.

## 5. Conclusions

A systematic study was performed to assess the safety implications of and offer guidance (see below) on tES stimulation in the presence of active and passive implants. Key safety considerations addressed in this study include localized E-field enhancement due to the presence of highly conductive contact, low-impedance capacitive pathways between contacts, and capacitive current injection into implant leads. While the focus was on tES, important aspects of the methodology, especially those related to the enhancement factors and the systematic exploration of active implant designs, can be leveraged for TMS and other forms of brain stimulation as well, upon revalidation of the approach. An important safety concern that still must be systematically investigated is the presence of burr holes through the skull with or without fixation screws, guidance channels, or similar.

### 5.1. Safety Guidance

Local exposure enhancements primarily concern brain tissue. They only minimally affect the already high fields near scalp electrodes, which are exposure limiting in terms of undesirable (confounding) sensations and/or the safety-relevant skin exposure quantities such as charge accumulation and current density. In terms of brain exposure safety during tES, undesired neuromodulation is the primary concern at low frequencies (sub-kHz), with E-field (magnitude and heterogeneity) constituting the limiting factor for brain exposure QoI [4]. In the presence of implants, the associated QoIs (i. e. E_ICNIRP,n_ and E_IEEE,n_) exhibit significant enhancement, such that highly localized stimulation of neural tissue near implanted electrodes under standard tES conditions cannot be excluded. The maximal E_ICNIRP,n_ and E_IEEE,n_ for DBS- or SEEG-like configurations can reach one order of magnitude (factor 10; see Table 1). The exposure scenarios associated with these enhancement resemble those of DBS in terms of focality and field gradient distribution. However, they are about 2-3 orders of magnitude lower than commonly employed DBS conditions (comparability is, however, limited by the different electrode implantation locations and frequency / pulse-shape). When interpreting tES impact results in the presence of conductive implants, proper consideration for that neuromodulatory effect is required, as some of the observed impact attributed to tES could be related to associated field enhancement (at locations identical or distinct from the stimulation target). This is of particular concern, when the implant serves as an electrophysiological signal acquisition electrode (e.g., SEEG), as the measurement is performed right where the confounding enhancement-related neuromodulation occurs.

In the absence of implants, temperature increase is typically not safety limiting at sub-kHz frequencies [4]. Therefore, the moderate increase (below 20 %, see Table 1) resulting from local field enhancement and power deposition in the presence of implants is of minor concern.

Elongated passive implants, such as stents made from conductive materials, can be associated with even larger enhancement factors than DBS and SEEG electrodes, such that their presence is often incompatible with tES.

The magnitude of capacitive effects are highly frequency dependent. At typical tES frequencies (100 Hz), capacitive coupling between contacts is negligible in comparison to the ohmic pathway through tissue. Capacitive coupling into the lead results in an injected current that generates local exposure upon flowing into tissue at implant contacts. When the lead is coiled around the tES electrode, such currents can give rise to exposures of comparable magnitude to those produced by (passive) local field enhancement, leading to similar concerns. However, in the case of tES in the kHz regime the situation changes, as the stimulation effectiveness decreases and the applied current strengths can strongly increase [4]: Brain temperature increase typically becomes the limiting factor, necessitating a reduction in stimulation power corresponding to the local-field-enhancement-related temperature increase (in addition to the above considerations about undesired stimulation). Capacitive coupling becomes much more relevant, which in combination with the increased stimulation current limit means that injected currents can increase by several orders of magnitude. For example, TIS at 100 kHz with 15 mA currents could in the worst-case result in capacitively injected currents with DBS-like stimulation magnitudes (milliamperes; the previous remark about the limited comparability in view of the different electrode implantation locations and frequency/ pulse-shape applies again).

To be conservative, additional safety margins might be required. The RMS difference of the enhancement factor prediction with the combined simulation reference (see Sec. 3.3) and the discretization error (see Sec. 3.1) suggest that a safety factor of 2 for field enhancement magnitudes is appropriate to achieve a coverage factor of *k* = 2 (95 %). On the other side, it is important to keep in mind that the determined maximal enhancement factors and capacitive coupling magnitudes already involve a series of worst-case assumptions regarding, e.g., electrode geometry, field alignment, and the lead-electrode proximity. Taking measures to avoid worst-case conditions, such as keeping lead loops away from stimulation electrodes, reducing implant exposure, and optimizing exposure angle, can reduce risks and avoid confounding neuromodulation. The data and approaches presented in this study can then be used to obtain less conservative enhancement and coupling estimates. Selecting less problematic implants (e.g., SEEG with shorter, well separated contacts when concurrent tES application is planned) can further reduce the safety concerns and increase the acceptable exposure magnitude, provided suitable alternatives exist.

Abandoned and leads with damaged insulations, especially in close vicinity of the tES electrodes, result in unwanted forms of stimulation and should constitute an exclusion criterion, when the risk cannot be bounded.

## Appendix A Complementary Results

**Figure A.1.**
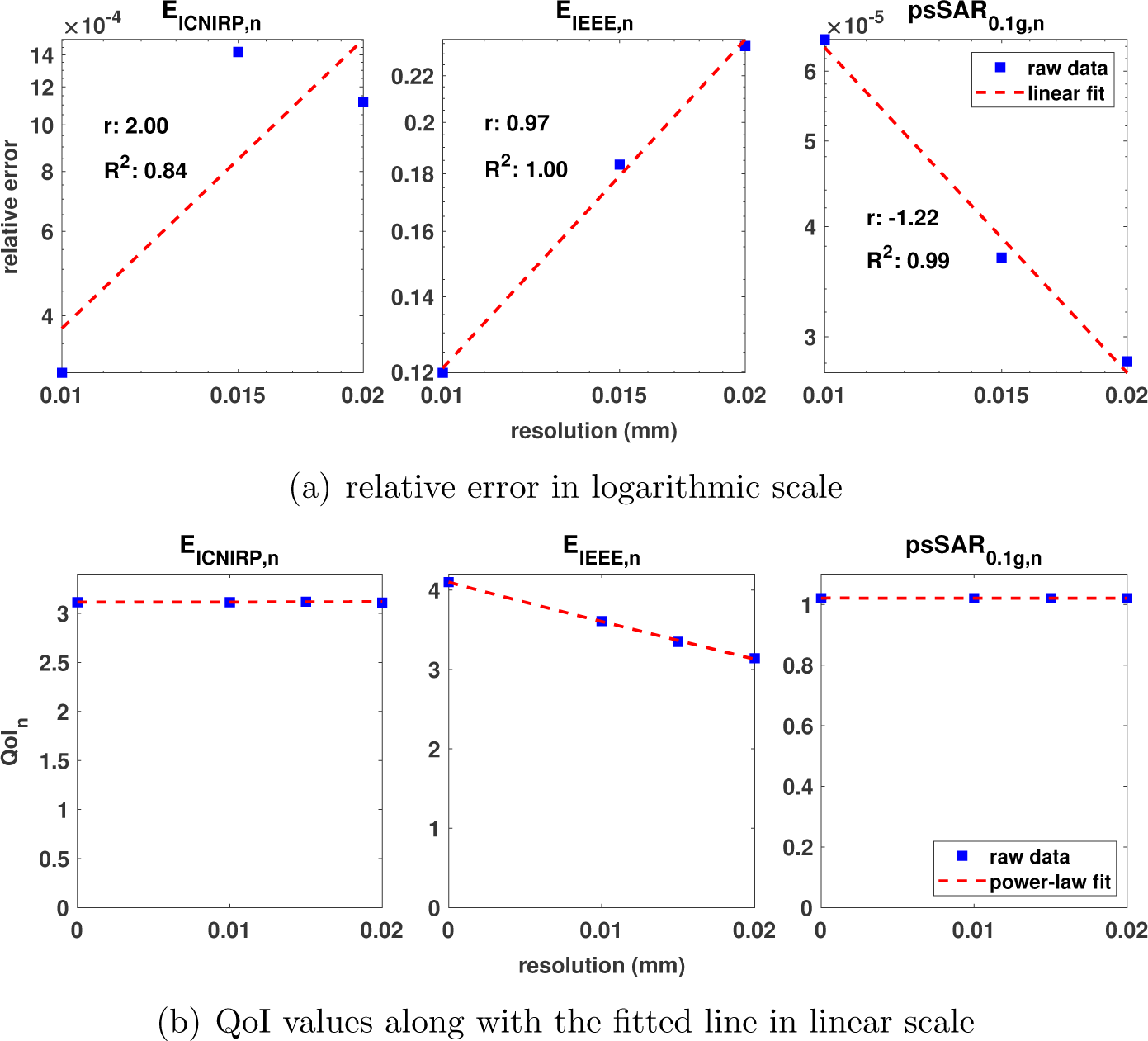
Discretization convergence analysis for the dosimetric QoIs E_ICNIRP,n_, E_IEEE,n_, and psSAR_0.1g,n_ (implant-parallel field incidence). (a) Log-log plots of relative discretization errors as a function of resolution; the linear regression slope is indicative of the order of the numerical scheme. (b) Power-law fit to quantity of interest (QoI) based on the linear regressions in the log-log plots. The interplay between these two views can be used to determine asymptotic QoI values at infinitely fine resolution (see Sec. 2.2.4).

**Figure A.2.**
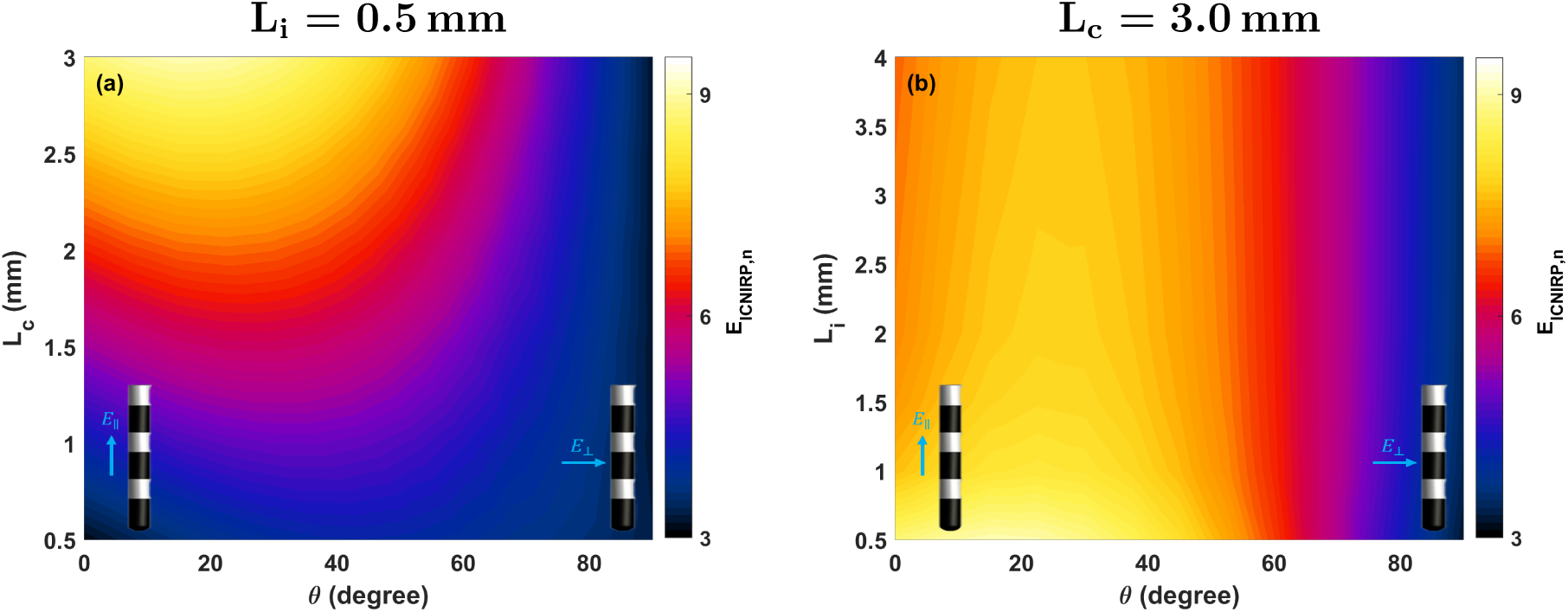
E_ICNIRP,n_ for the 0.8 mm diameter implant as a function of field-incidence angle and (a) contact length (fixed insulator length *L_i_*=0.5 mm) and (b) the insulator length (fixed contact length *L_c_*=3.0 mm).

**Figure A.3.**
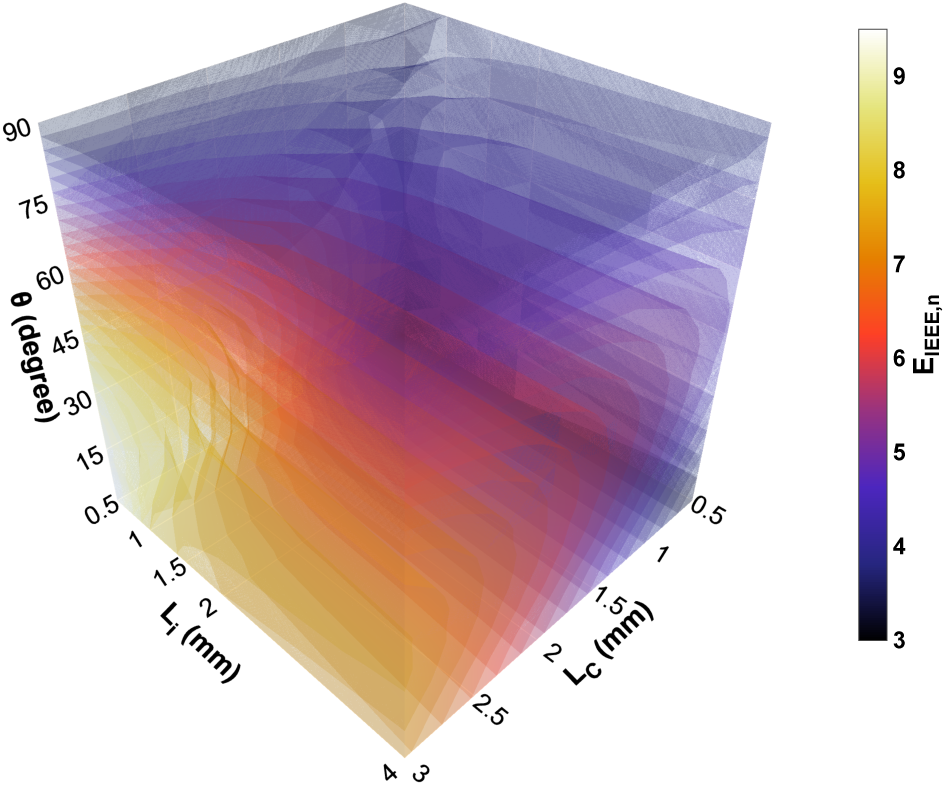
E_IEEE,n_ for a 0.8 mm diameter implant as a function of field-incidence angle, contact length, and the insulator length.

**Figure A.4.**
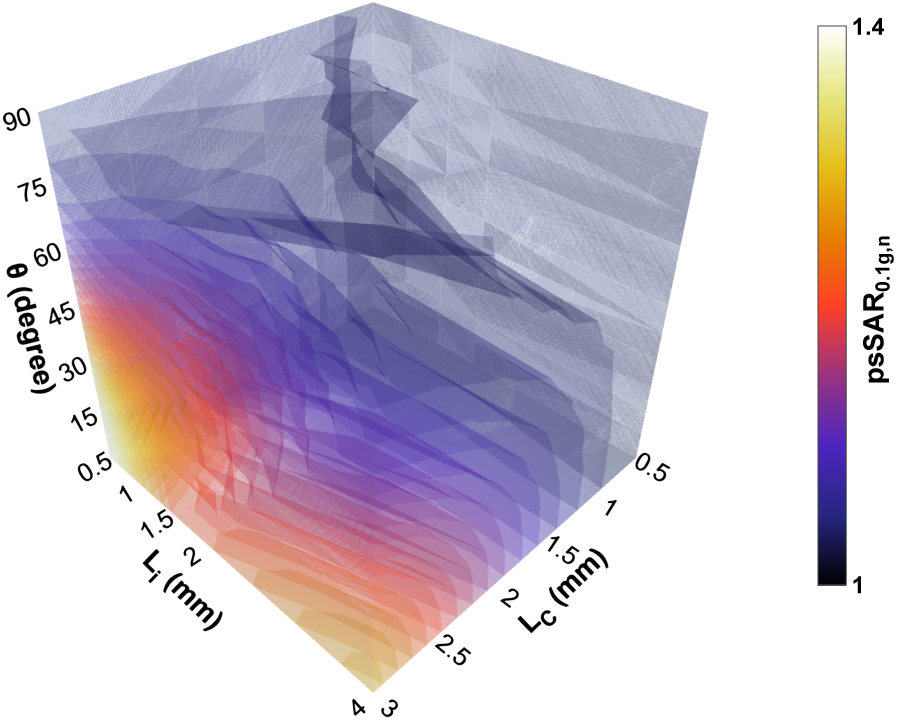
psSAR_0.1g,n_ for a 0.8 mm diameter implant as a function of field-incidence angle, contact length, and the insulator length.

**Figure A.5.**
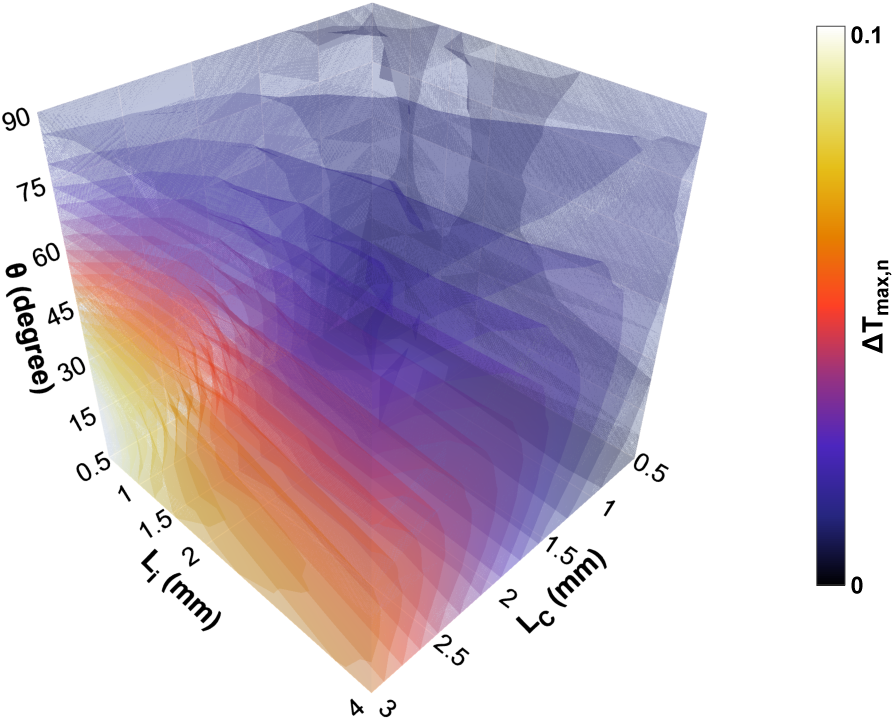
ΔT_max,n_ for a 0.8 mm diameter implant as a function of field-incidence angle, contact length, and the insulator length.

**Figure A.6.**
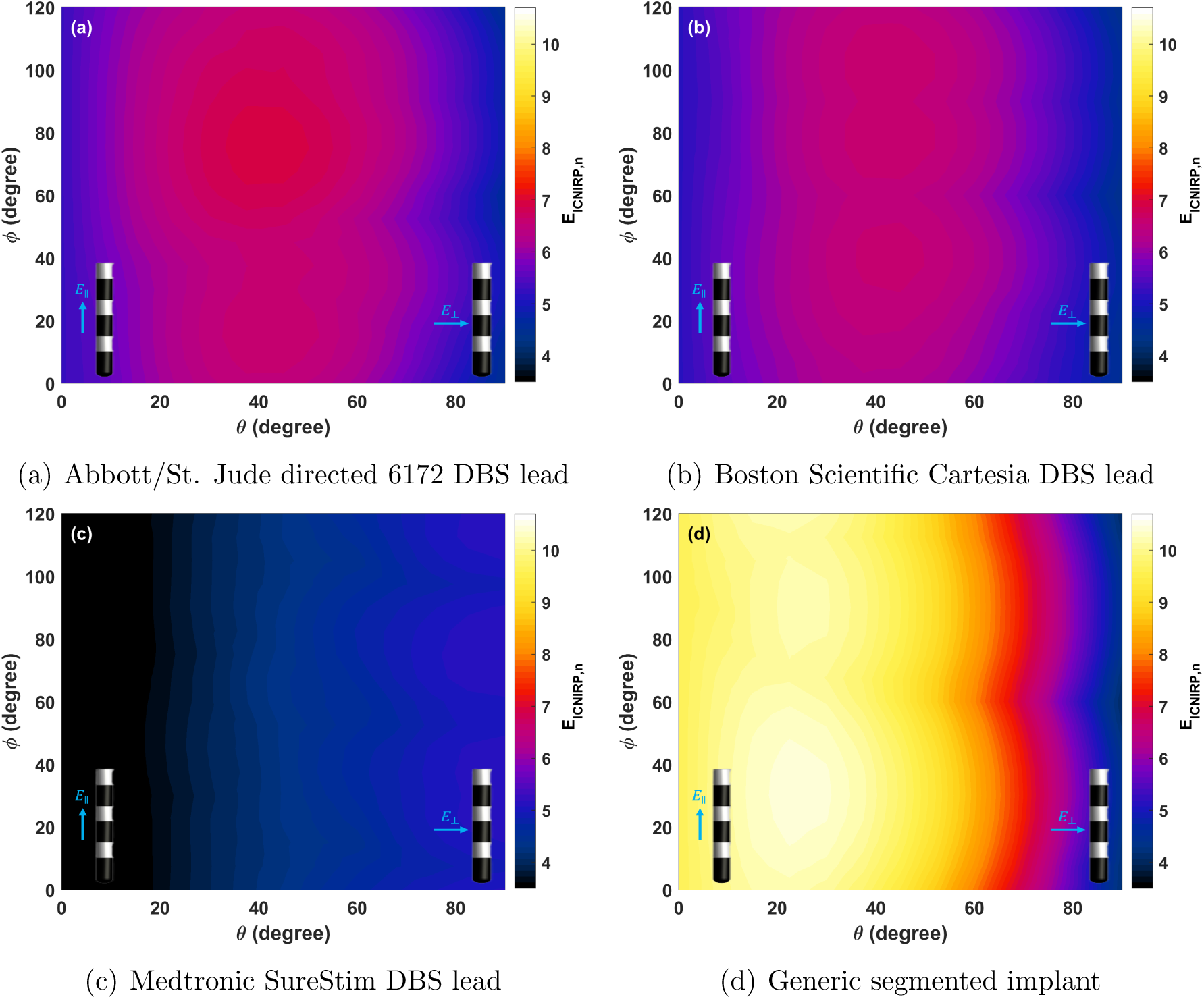
E_ICNIRP,n_ for the segmented and HD leads (see Fig.3(e-h)) as a function of field-incidence angles. The generic segmented implant (d) is associated with the worst case of all generic implants (*d*=1.4 mm, *L_c_*=3.0 mm, and *L_i_*=0.5 mm).

**Figure A.7.**
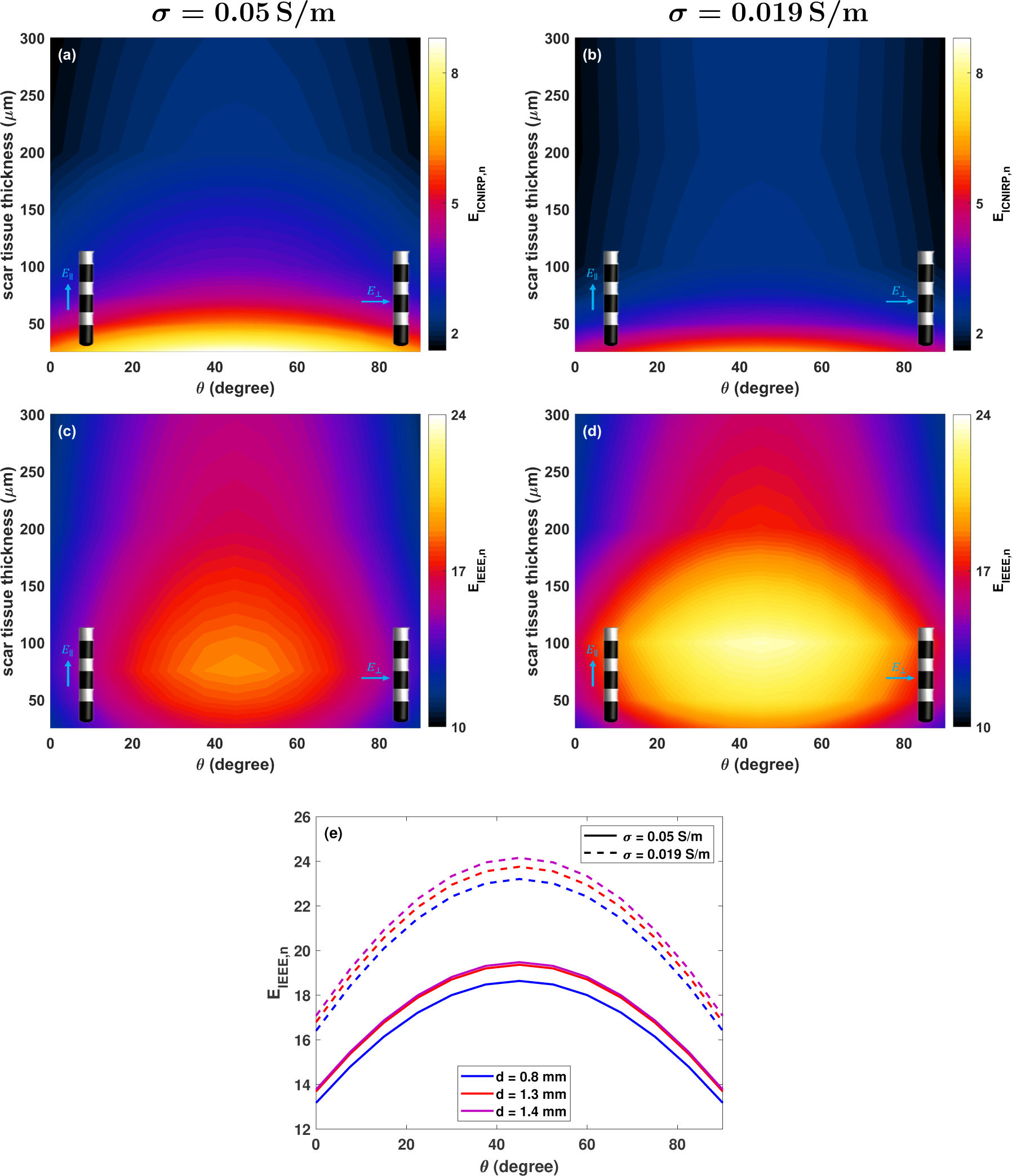
(a,b) E_ICNIRP,n_ for the worst case of generic implant with 0.8 mm diameter (*L_c_*=3.0 mm and *L_i_*=0.5,mm) leads encapsulated in scar tissue (Fig.3(j)) as a function of field-incidence angles and scar tissue thickness. (c,d) E_IEEE,n_ for the worst case of generic implant with 0.8 mm diameter (*L_c_*=3.0 mm and *L_i_*=0.5,mm) leads encapsulated in scar tissue (Fig.3(j)) as a function of field-incidence angles and scar tissue thickness. The E-field in scar tissue is considered in this analysis. (e) Maximum E_IEEE,n_ for worst case generic implant (*L_c_*=3.0 mm, *L_i_*= 0.5 mm) for 100 *µ*m scar tissue thickness as a function of the field-incidence angle. The E-field in scar tissue is considered in this analysis.

**Figure A.8.**
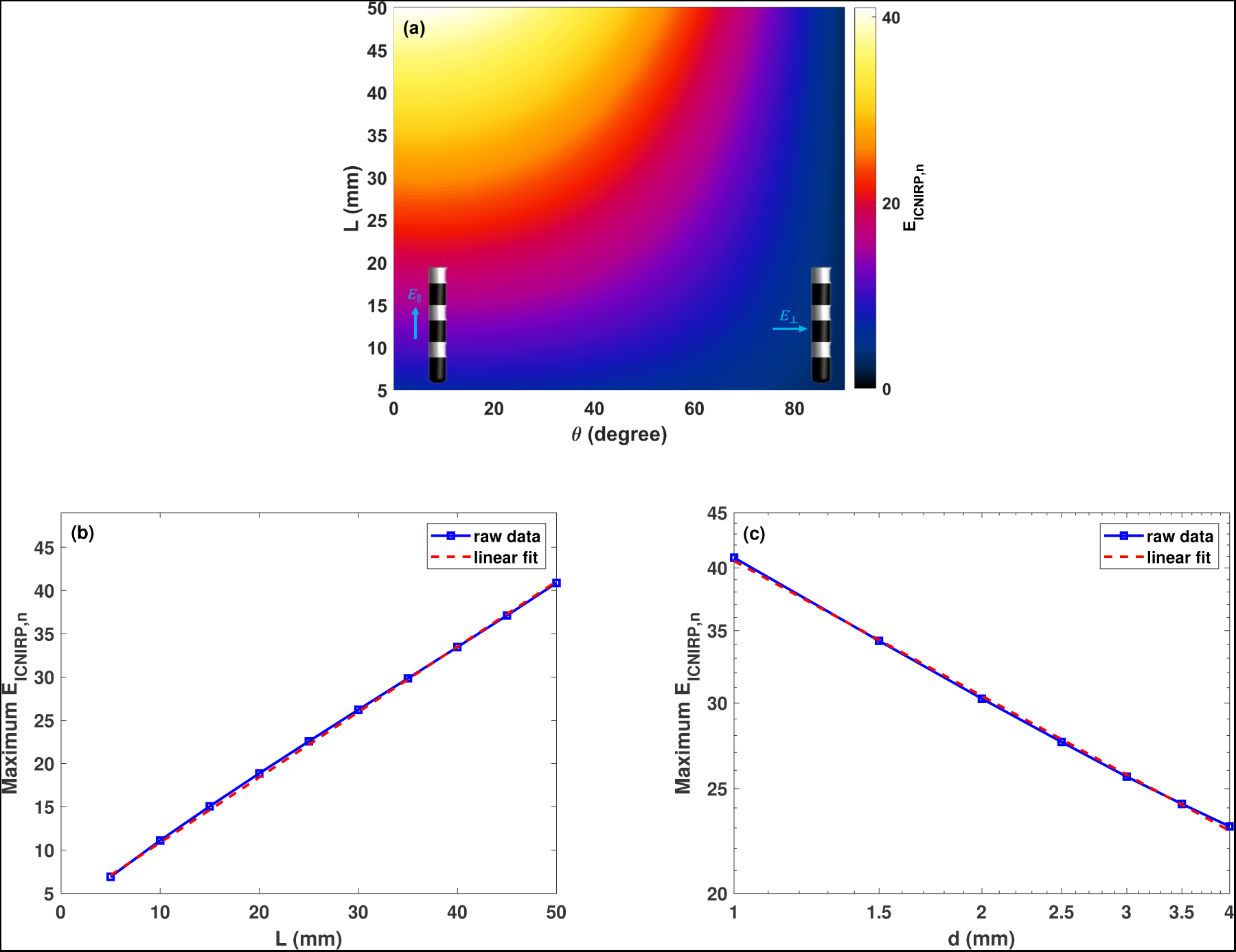
(a) Maximum E_ICNIRP,n_ for an elongated implant with 1 mm diameter as a function of the implant’s length and the field-incidence angle. Maximum E_ICNIRP,n_ for elongated implant (b) as a function of implant’s length when the implant’s diameter is 1 mm (maximum E_ICNIRP,n_ *≈* 0.8 *mm^−^*^1^*L* + 3.3 where L is in mm), and (c) as a function of implant’s diameter when the implant’s length is 50 mm (E_ICNIRP,n_ *≈* 40.60 *mm*^0.41^*d^−^*^0.41^, where d is in mm). Note the logarithmic *x* - and *y* -axis scales in (c).

**Figure A.9.**
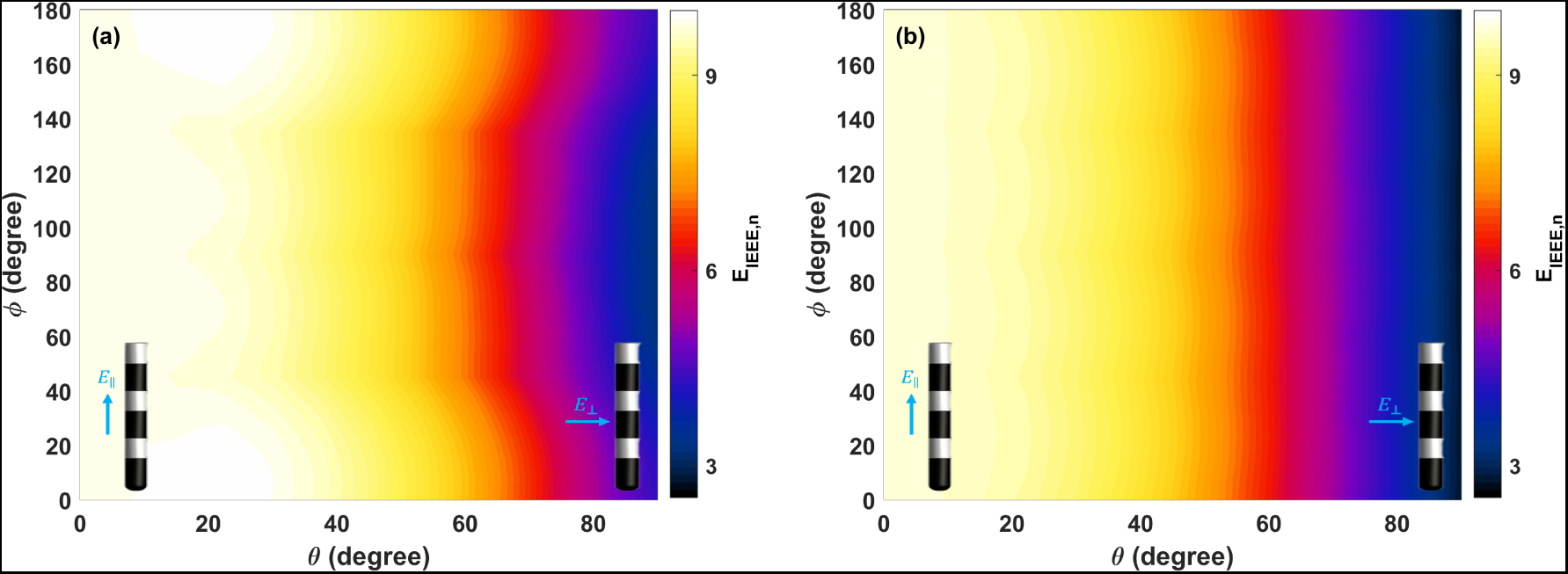
E_IEEE,n_ for the worst case of the generic implant with 0.8 mm diameter (*L_c_*=3.0 mm and *L_i_*=0.5,mm) leads by considering anisotropic conductivity for tissue simulating medium. The principal tensor axis was assigned the white matter axial conductivity value from the IT’IS tissue properties database [56], while the radial value was assigned to the two perpendicular directions. That principal axis was (a) perpendicular to the the implant and (b) aligned with it. Anisotropy increases the maximum E_IEEE,n_ from 9.4 to 10.0.

